# Simulating Neutron Protein Crystallography Experiments: Applications to the Development of the NMX Instrument at ESS

**DOI:** 10.64898/2026.03.26.714568

**Authors:** Mads Bertelsen, Peter K. Willendrup, Sunyoung Yoo, Adriano Meligrana, David McDonagh, Justin Bergmann, Esko Oksanen, Aaron D. Finke

**Affiliations:** European Spallation Source ERIC, Data Management and Scientific Computing, Kongens Lyngby, Denmark; European Spallation Source ERIC, Lund, Sweden; Department of Computer, Control, and Management Engineering, Sapienza University of Rome, Italy; STFC Rutherford Appleton Laboratory, Didcot, UK; LINXS - Institute of Advanced Neutron and X-ray Science, Lund, Sweden; Division of Computational Chemistry, Lund University, Lund, Sweden

**Keywords:** Neutron macromolecular crystallography, Monte Carlo simulations, Neutron diffraction

## Abstract

Monte Carlo neutron ray-tracing simulations of time-of-flight (TOF)-Laue neutron macromolecular crystal diffraction (*n*-MX) using the McStas software package were done for the upcoming NMX Macromolecular Diffractometer at the European Spallation Source. Splitting neutron rays that arrive at the crystal lead to dramatic improvements in event formation with minimal computational overhead. The simulated event probability data was sampled using a new single-pass weighted reservoir sampling method, and processed like real *n*-MX data using DIALS. The effects of air and beamstop scatter on simulated data was investigated.

**Synopsis:** Monte Carlo simulations of neutron protein diffraction experiments provide useful data that models instrumental components that interact with neutrons, as well as the crystal diffraction itself. These data can be applied to instrument development, such as the commissioning of the NMX Macromolecular Diffractometer at ESS.

## 1 Introduction

The work detailed herein answers the question: how can one perform a neutron experiment without neutrons? In the strictest, most mundane terms, it is impossible; but with a computer and a sophisticated enough understanding of the underlying physics, we can certainly *model* such an experiment with a precision that yields useful insight. Thus, the current standard is to make use of the computational tools that are good at modelling experiments, thereby pushing the boundaries of those experiments. Such synergy is at the core of this work.

### 1.1 The NMX Instrument at ESS

The European Spallation Source (ESS) is a European project to build what will become the world’s most powerful neutron spallation facility.^1^.(Peggs, 2013; Garoby *et al*., 2017) The NMX Macromolecular Diffractometer,(Markó *et al*., 2020) — the ESS’s flagship instrument for neutron macro-molecular crystallography (*n*-MX) — is designed to benefit from the long neutron pulse and the bright moderators, which are the unique features of the ESS.

The neutron scattering factors of hydrogen and deuterium are on par with those of carbon and oxygen, leading to accurate and refineable H-atom positions from *n*-MX data(Moody, 2020; Meilleur *et al*., 2006). More commonly used methods such as X-ray MX, electron MX, and cryo-electron microscopy do not provide precise hydrogen atom positions, which makes *n*-MX a truly complementary technique to these more widely available methods. However, *n*-MX remains experimentally challenging, so it has thus far been used on a limited number of macromolecules. The ambition at ESS is to expand the utility of *n*-MX to a broader range of biologically relevant systems. (Oksanen, 2026)

The “entry barrier” to *n*-MX is having crystals that are not only large (on the order of 1 mm^3^),(Budayova-Spano *et al*., 2020) but also isotopically-labelled, with the hydrogens replaced with deuterium to eliminate the strong incoherent neutron scattering from ^1^H (though it is often sufficient to simply swap labile protons with deuterium in D_2_O). Higher neutron flux, such as what the ESS will provide, can greatly lower the barrier to entry by requiring smaller crystals. Since neutron sources will always have lower flux than X-ray sources, we want to take advantage of every neutron that is available. Thus, Laue diffraction techniques, which use a wider bandwidth of radiation, improves neutron economy;(Niimura *et al*., 1997) and time-of-flight Laue (TOF-Laue) techniques, which exploit the de Broglie wave nature of neutrons, are unique to neutron crystallography and can improve signal-to-noise and facilitate refinement.(Bull *et al*., 2014; Peters & Jauch, 2002)

NMX is also designed to overcome challenges in the *n*-MX experiment unrelated to the neutron source. Most *n*-MX instruments have detectors at fixed geometries, in order to maximize angular coverage at the cost of limiting the resolvable unit cell size.(Schultz *et al*., 2005; Keen *et al*., 2006; Tanaka *et al*., 2009) NMX, by contrast, will feature a variable-geometry detector setup to allow resolution of reflections from crystals with large unit cells, *i*.*e*. by increasing the sample-to-detector distance. Trading solid angle coverage per exposure to achieve this will nonetheless expand the *n*-MX technique to broaden the range of systems that can be studied.

High-quality simulation data allows to test our ability to translate the promise of these innovations to real data well before the instrument opens. Fortunately, much of the groundwork has already been laid for simulating *n*-MX data.

### 1.2 Simulations for Neutron Instrument Planning

There are a few software packages available for neutron simulations, including (but not limited to) McStas, VITESS,(Manoshin *et al*., 2011) and Geant4.(Allison *et al*., 2016) The former, McStas, is used in this work. McStas is an open-source software package for Monte Carlo (MC)-based raytracing of slow neutrons and their interaction with matter.(Willendrup *et al*., 2014; Willendrup & Lefmann, 2020; Willendrup & Lefmann, 2021) McStas has been used extensively for simulations of experiments at both current and planned neutron instruments.(Udby *et al*., 2011) It has been used to validate the design of every instrument at ESS so far, including NMX.(Toft-Petersen *et al*., 2025; Markó *et al*., 2020)

A complete description of the underlying algorithms driving McStas is beyond the scope of this work, so we will dispense with a simplistic picture. We recommend consulting the McStas documentation for a complete description. In McStas, a neutron is treated as a *ray* with an associated *probability*. The properties and trajectory of each ray are defined by three things: the **source**, which define the ray’s initial properties; the **instrument**, a list of all components the ray can interact with, and the **monitors**, which record the results. Each component in the instrument will have a calculated probability of interacting with a ray, modifying its momentum vector. If a ray crosses a monitor— in our case, an ideal 2-dimensional area detector made up of pixels— it is recorded in so-called “event mode,” which records the ray’s time of arrival (TOA), energy,^2^ and total probability.

The computational logic behind McStas is simple but powerful. The instrument file containing the experimental parameters and components is converted into C-code, which can be compiled by any C compiler such as *gcc* or *clang*. The simulations themselves are done by simply running the resulting binary. McStas requires few external libraries, but parallelization— a necessity for this work— requires the *MPI* library. Because each simulation is that of an uncorrelated neutron ray trajectory, McStas is highly parallelizable and is well-suited to high-performance computing (HPC).

### 1.3 Simulation of Neutron MX Experiments

As mentioned above, McStas has considerable utility in neutron instrument design, particularly for neutron optics. It was also used in the optics design for NMX(Markó *et al*., 2020). McStas can be used to do realistic simulations of single-crystal diffraction experiments, but there are some important challenges that need to be addressed in order to properly simulate *n*-MX data:

- Sampling all of reciprocal space is difficult, as there are a large number of reflections that must be sampled, which can be *>*10^6^.
- Since MC simulations also model the probability of a protein crystal scattering a neutron ray, and such probabilities are low, a large number of simulations are often necessary to fully cover reciprocal space.
- As a result of the above two issues, computation— in terms of processing power and memory— can be costly.

For NMX specifically, an additional difficulty is doing simulations for an instrument under construction that has no real data to compare it against (yet) and no comparable instrument exists elsewhere. The simulated data must nevertheless be realistic, and so should be processed the same as any real *n*-MX data from a real instrument. That is also the benchmark we will use.

A primary inspiration for simulating *n*-MX experiments is MLFSOM(Holton *et al*., 2014), a program developed to generate X-ray MX data from a list of structure factors and as exhaustive a list as possible of experimental parameters. The primary goals of MLFSOM were to better understand the sources of experimental error in MX experiments. Our current aims are different: to demonstrate the feasibility of using MC simulations for *n*-MX experiments, and applying them to NMX instrument commissioning. Ultimately, however, our approach should also be applicable for modelling sources of experimental error in *n*-MX experiments.

## 2 Setup of Computations with McStas

To perform *n*-MX simulations with McStas, one needs a complete description of the instrument to be modelled and a list of Bragg reflection data.

### 2.1 The Instrument File

The McStas instrument file contains all the component information for the instrument, from source to detector. Example instrument files used for NMX are in the Supporting Information.

#### 2.1.1 Neutron Source and Moderator

McStas itself does not model neutron sources (e.g. fission reactors or spallation sources) or the interaction of sources with neutron moderators; these can be done with other simulation programs such as GEANT4 and MCNP(Kulesza *et al*., 2024). Instead, it provides the distribution and geometry of the moderator output calculated elsewhere. The neutron *moderator*, not the spallation target, is considered the “source” of the neutrons for our simulations.

For the NMX instrument, we use the ESS “butterfly” moderator component,(Andersen *et al*., 2020). One can specify the relevant beam port for this moderator in the instrument file; for NMX, this is beamport W1. The ESS has a long pulse length compared to other spallation sources (2.86 ms, compared to 0.2-0.5 ms elsewhere), leading to a significant uncertainty to the calculated TOF. For this reason, the sample position of NMX is quite far from the source (157 m) to improve TOF resolution.(Markó *et al*., 2020) TOA uncertainty is propagated in neutron ray-tracing simulations.

#### 2.1.2 Neutron guides

The NMX beam delivery system is designed to deliver to the sample a maximum 5×5 mm^2^ beam of neutrons with ±0.2° divergence. The neutron guides are optimised for wavelengths *>*1.8 Å and all the optical components from the moderator to the experimental cave are modelled in McStas.

#### 2.1.3 Choppers

NMX uses two wavelength-defining disk choppers to provide a typical wavelength range of 1.8 to 3.55 Å or 2.2 to 3.95 Å. (Markó *et al*., 2020) In McStas, one can specify the energy range of neutrons coming out of the moderator to improve calculation efficiency. In tandem, the phase of the choppers is calculated for the specified wavelength range. We can use this to simulate out-of-phase errors and deviations from ideal behaviour, including jitter effects.

#### 2.1.4 Collimation system

The collimating devices in the NMX cave, including slits and pinhole apertures, define the beam size and divergence at sample. In addition, the effects of air scatter on the neutron beam must also be modelled from the last vacuum window as the last parts of the system are in air. We use the Union component system in McStas to define components of complex geometry,(Bertelsen, 2017) and that are made of materials whose neutron interaction properties have been modelled using the NCrystal library.(Cai & Kittelmann, 2020) Using the Union system enables multiple scattering between a complex set of component geometries. For example, the NMX pinhole is made of Gd, and the full geometry of the pinhole system, along with the surrounding air, is defined as a Union component.

#### 2.1.5 Sample

The Single crystal component in McStas as used in NMX simulations is described in greater detail in section 2.2.1.

#### 2.1.6 Beamstop

For the initial simulations, a simple, idealistic beamstop concept was used, wherein any neutron ray that arrived at the crystal but did not scatter was discarded at that point. These simulations have very little background from air scattering and so are “unrealistic”, but nonetheless useful for verifying the quality of the simulations up to the crystal. Later, more complex scenarios were simulated. First, we included a “perfect” beamstop component— a disc of 1 cm diameter that absorbed any neutron that crossed it— set 1 cm downstream from the sample, in order to probe the effects of air scattering after the crystal (not shown). Ultimately, a Union beamstop of radius 3.5 mm made of ^6^LiF powder in a 1 mm thick Al casing, set 1 cm downstream of the sample, was used as the prototype NMX beamstop. As the beamstop component and surrounding environment become more complex, the number of recorded events increases, as well as the computation time.

#### 2.1.7 Detectors

The NMX detector system (Pfeiffer *et al*., 2016; Pfeiffer *et al*., 2026) is quite unique, in that it consists of three fully mobile, wide-area triple-gas electron multiplier (GEM) panels, in contrast to the fixed detector geometry of most other *n*-MX TOF-Laue instruments. The three detector panels are mounted on mobile robot arms with nearly complete coverage of the top hemisphere around the sample. To avoid collisions, the detector positioning system of NMX uses predetermined detector configurations. The current setup has 11 configurations, but based on simulations and commissioning experiments these can be modified and other configurations added.

In McStas, each detector panel component is modelled as a 1280×1280 grid of pixels of size 400*µm*. The detectors themselves were modelled as “ideal” monitors with perfect detection efficiency and no point spread function. Simulations of neutron detectors can be done in a limited manner in McStas, which we will explore in the future; more sophisticated GEM detector simulations have been described elsewhere.(Pfeiffer *et al*., 2019) The detector box components were not modelled for these simulations.

Neutron monitor components in McStas can be configured to “restore” neutron rays that hit them, effectively making the monitor transparent. All 33 planned detector positions can thus be simulated from a single instrument file when the `restore_neutron` option is turned on for each monitor component. Strictly speaking, this is not the same as simulating the 11 detector configurations independently, due to the fact that a neutron ray only interacts with the first monitor exit volume it crosses and ceases to interact with air after crossing it (but will still be measured by any other monitor it crosses). Nonetheless, the differences between simulations run with one configuration at a time and simulations with all configurations run simultaneously are small, and simulations run with all configurations is ten times more efficient in CPU time.

### 2.2 Simulating a Protein Crystal

The simulated protein crystal is defined by two parts: the Single Crystal component in McStas and the list of Bragg reflections.

#### 2.2.1 The Single crystal component

The Single crystal component in McStas is defined by a list of Bragg reflections, unit cell axes, crystal size, lattice plane spacing uncertainty Δ*d/d*, and mosaicity *η*. For our calculations, Δ*d/d* and mosaicity were kept low, as they tend to be low for protein crystals at room temperature. Crystal volumes of 1 mm^3^ were used. The crystal orientation was set so the (100), (010), and (001) planes were perpendicular to the McStas X, Y, and Z axes, respectively. Solvent surrounding the crystal and quartz capillary crystal holders, both of which are present in real *n*-MX measurements, were not modelled for these simulations.

The scattering process for a Single crystal component works as follows. A neutron ray of a particular energy and vector interacts with the component, and its probability of interaction is calculated. If the neutron ray interacts with the crystal, it is SPLIT (see below) and reflections that fulfil the Bragg condition are taken from the complete list (Section 2.2.2). The list of reflections chosen is effectively a “shell” with the same radius as the Ewald sphere, with a shell thickness of 3*σ* deviation. For more details, see §8.4 of the McStas component manual. Scattering events were limited to 0th and 1st-order scattering only, corresponding to transmission through the crystal and a single scattering event from the crystal, respectively. Including second-order scattering or higher increases computation time by an order of magnitude, and such events have negligible probabilities and so can be ignored.

One of the biggest sources of computational inefficiency in McStas is the scattering of neutron rays from the crystal. The probability of a neutron scattering coherently off a protein crystal is already quite low. To improve efficiency, the SPLIT instruction in McStas is used for the Single crystal component. In brief, a neutron ray that hits the crystal and scatters is “split” (best thought of as copy/pasting the neutron ray) into a defined SPLIT number of neutron rays, which will scatter off the crystal as well. Since the absolute number of neutron rays simulated relative to the total number of scattered neutron rays calculated is irrelevant, we can set the SPLIT value to a high number to greatly improve computation efficiency. For example, 1e12 neutron simulations without SPLIT leads to ≈6.8e5 event probabilities recorded; with SPLIT 11000, that number jumps to ≈2.3e10 events (Table 1), with a tenfold increase in SPLIT leading to a tenfold increase in recorded event probabilities.

**Table 1:** Effect of *N*_*simulations*_ on recorded *N*_*events*_.

| $N_{simulations}$ | $N_{events}$ | time<sup>a</sup> |
| --- | --- | --- |
| 5e9 | 3.81e7 | 21 s |
| 5e10 | 3.79e8 | 3.75 min |
| 5e11 | 3.77e9 | 37.8 min |
| 5e12 | 3.77e10 | 6.04 h |
| 1e12 (no SPLIT) | 6.84e5 | 1.25 h |

<sup>a</sup> Using 1280 MPI processes

Incoherent scattering from the crystal can be significant, particularly if the hydrogen (^1^H) content is high. In this work, we are only considering perdeuterated crystals in D_2_O, and thus the total incoherent scattering is low. Absorption, coherent scattering, and incoherent scattering cross-sections of protein crystals are user-provided and included in the list of Bragg reflections (see below). They can be easily calculated from the molecular formula of the protein and knowledge of the neutron scattering and absorption cross-sections; the *PeriodicTable* Python library(noa, 2026) nicely automates the process (see the Supporting Information).

#### 2.2.2 List of Bragg Reflections

McStas must be provided with a complete list of Bragg reflections, defined by a Miller index (*h, k, l*) and structure factor F^2^ in barns. Currently, McStas does not calculate or know anything about crystal symmetry and assumes the space group is *P* 1. Critically, it also does not calculate Friedel pairs. When preparing a list of reflections for McStas calculations, it is thus necessary that a *full sphere* of reflections to a given resolution is provided, not just the hemisphere one would get from taking a list of reflections with symmetry and expanding it to *P* 1 in most programs. *Any reflection not on the list will not be modelled*.

There are a number of ways to calculate structure factors of protein crystals with neutrons. The open-source structural biology toolkit Gemmi(Wojdyr, 2022) is excellent for this, as is *phenix*.*fmodel* from the Phenix suite of MX software.(Liebschner *et al*., 2019) The latter tool has the advantage of modelling coherent solvent scattering automatically. Modelling solvent can also be done with Gemmi in a few steps. However, do note that both methods require manually adding Friedel pairs to the generated lists. Using Gemmi with Pandas in Python can easily perform this operation.

Reflection lists for perdeuterated Pyrococcus furiosus rubredoxin (PDB 4AR4)(Cuypers *et al*., 2013) were used for this work. Experimental neutron data with refined D positions were used as a starting point. A complete list of neutron scattering factors to either 1.2 or 0.6 Å was calculated from the deposited coordinates and refined against the experimental structure factor data to account for solvation. The Jupyter notebook used to generate the .lau reflection files with Gemmi is provided in the Supporting Information.

### 2.3 Hardware requirements

McStas output binaries use MPI for highly parallelized computations; and with the SPLIT instruction, fewer simulations from source to sample are required. Simpler simulations, such as those without realistic beamstops and without large volumes of air, require fewer total ray-tracing simulations than those that do have those things. For the NMX instrument, a “simple” simulation of rubredoxin crystal, with an ideal beamstop that simply discards any neutron that does not scatter off the crystal, requires a minimum of ca. 1e10 simulations, and up to ca. 5e11 simulations for more complex ones (e.g. modelling the scattering from a realistic beamstop, see below).

We make heavy use of HPC resources for these calculations, in particular using the LUMI supercomputing cluster in Kajaani, Finland. A simple simulation with *N*_*simulations*_ = 5*e*11 and *SPLIT* = 11000 takes approximately 30 minutes on LUMI using 1280 CPUs, and processing time scales roughly linearly with the number of CPUs used. While it is always prudent to exploit the best possible resources one can access, we stress HPC resources are *not* necessary to perform these calculations. One can easily do these simulations on a higher-end, consumer-grade system if it has sufficient memory^3^— they will just take hours to days instead of minutes. Storage can be a practical issue, as the output files for these simulations can run into the hundreds of gigabytes to terabytes. However, the output of multiple McStas runs can be combined easily, so one can offload the results of smaller runs onto offline storage if needed, and combine later during processing.

Parallelization for this work is entirely done with MPI. We are currently working on improving GPU acceleration for parallel computations in McStas. Currently, McStas has limited GPU support for NVIDIA GPUs but it is not optimized for SPLIT Single crystal components.

## 3 Converting Event Probabilities into Events

When the McStas monitor components (the detectors) record in event mode, as they do here, the recorded event is saved as an array with the neutron’s TOA, monitor pixel location, and the probability of the neutron trajectory. McStas only calculates *probabilities* of neutron events, not actual events. To convert the list of event probabilities *p* into a list of events *N*, we must sample the list of probabilities in full, treating each event as independent. The best solution to this is *weighted reservoir sampling with replacement*.(Park *et al*., 2007; Shekelyan *et al*., 2023)

Therein lies a problem: in order to fully sample reciprocal space for protein crystals in our *n*-MX simulations, we must record billions of events, totalling hundreds of gigabytes, if not terabytes, of data.^4^ Weighted reservoir sampling exists in statistical analysis software packages such as Numpy, but those methods use multi-pass algorithms that require the full list of event probabilities to be loaded into memory first. This is impossible in our case.

Instead, we rely on an algorithm for *single-pass* weighted reservoir sampling.(Meligrana & Fazzone, 2026) This algorithm generates an event list of size *N* — the reservoir — from a list of probabilities *p*, but only requires reading each element once; in other words, *p* need not be fully loaded into memory. This algorithm has been proven to generate equivalent reservoirs as multi-pass algorithms. The method also allows for random sampling from *streamed* data, that is, it can update the reservoir in real-time as elements in *p* arrive, providing access to simulating streamed instrument data.

Our *stream sampling mcstas* software uses the StreamSampling.jl library(Meligrana, 2026), which itself is based on the OnlineStats library(Day & Zhou, 2020) in the Julia language(Bezanson *et al*., 2017), to read a list of McStas event probabilities and generate a list of events with pixel location and TOA, much like a real neutron detector would.(Finke, 2026) In this way we can generate “real” event data from the McStas event probability data, in the same format (NeXus TOFRaw) that will be output when NMX comes online.

Another method that can be used to convert probabilities into events is to simply multiply each event probability by a large integer constant, thus converting each probability measurement into integer detector “counts”. These counts can then be binned and histogrammed. Doing this effectively simulates collecting an infinitely-long dataset, as the event list *N* should look like *p* as *N* → ∞. We have opted for weighted random sampling in this work, as we do not plan on collecting infinitely-long datasets.

## 4 Data Reduction and Analysis

### 4.1 Binning and histogramming

Neutron TOF-Laue data must be binned and histogrammed by TOF in order to be processed as “monochromatic” crystallographic data. We have developed the ESSNMX software package to process event data from not just NMX, but also our simulated data. It is based on the *scipp* package,(Heybrock *et al*., 2020) which provides numeric arrays with labels and unit measurements. ESSNMX converts measured TOA to TOF, and outputs the binned/histogrammed event data in the NeXus NXLaueTOF format. This is the same format that NMX will use when it comes online. ESSNMX also computes pixel positions for each detector based on the NXtransformation information in the TOFRaw format; NXtransformation is calculated from the McStas geometry in *stream sampling mcstas*.

### 4.2 Spotfinding, Indexing, and Integration

The latest version of the DIALS software package (v3.27.0)(Winter *et al*., 2018) includes support for spotfinding, indexing, and integration of *n*-MX TOF-Laue data from various instruments, as well as the simulated data from this work. DIALS imports the binned/histogrammed event/TOF data from ESSNMX. Spotfinding, indexing, and integration are done on the binned images. The indexing step is particularly useful for instrument commissioning, as we can test the sensitivity of measured detector geometry on indexing. The NMX detectors will be fully mobile in three dimensions, so reproducible positioning and accurate measurement of detector positions will be critical parameters for reliable indexing of NMX data.

Integration is done with the `dials.tof_integrate` routine now available in DIALS. A number of reflection profiling algorithms are available, including simple summation (which does no profiling), so-called “seed skewness” integration,(Peters, 2003) 1-D line profiling,(Tomoyori *et al*., 2013; Yano *et al*., 2016) and 3-D surface profiling.(Gutmann, 2017) The latter two methods tend to give the best I/*σ*(I) for the simulated reflections, at the cost of (slightly) higher computation time.

The Mantid software package(Arnold *et al*., 2014) can also be used to bin/histogram and reduce the simulated *n*-MX data. McStas can output the Mantid-style instrument data in XML format needed to load the sampled data into Mantid. Both software packages will give similar results.

## 5 Results and Discussion

### 5.1 Modelling Bragg Diffraction

Simple simulations of rubredoxin with NMX “box” detector geometry were performed with an ideal beamstop (i.e. any neutron not scattered by the crystal was discarded at that point). Fig. 1 shows a histogram of the sum of event weight probabilities by pixel, and the corresponding sampled dataset. As it is a Monte Carlo simulation, the number of simulated rays determines the completeness of the simulation space around the detectors. Table 1 Shows the effect of *N*_*simulations*_ on the number of events recorded on the three detectors, *N*_*events*_. All other things being equal, a factor of 10 increase in *N*_*simulations*_ leads to a tenfold increase in events measured on the detectors.

**Figure 1.**
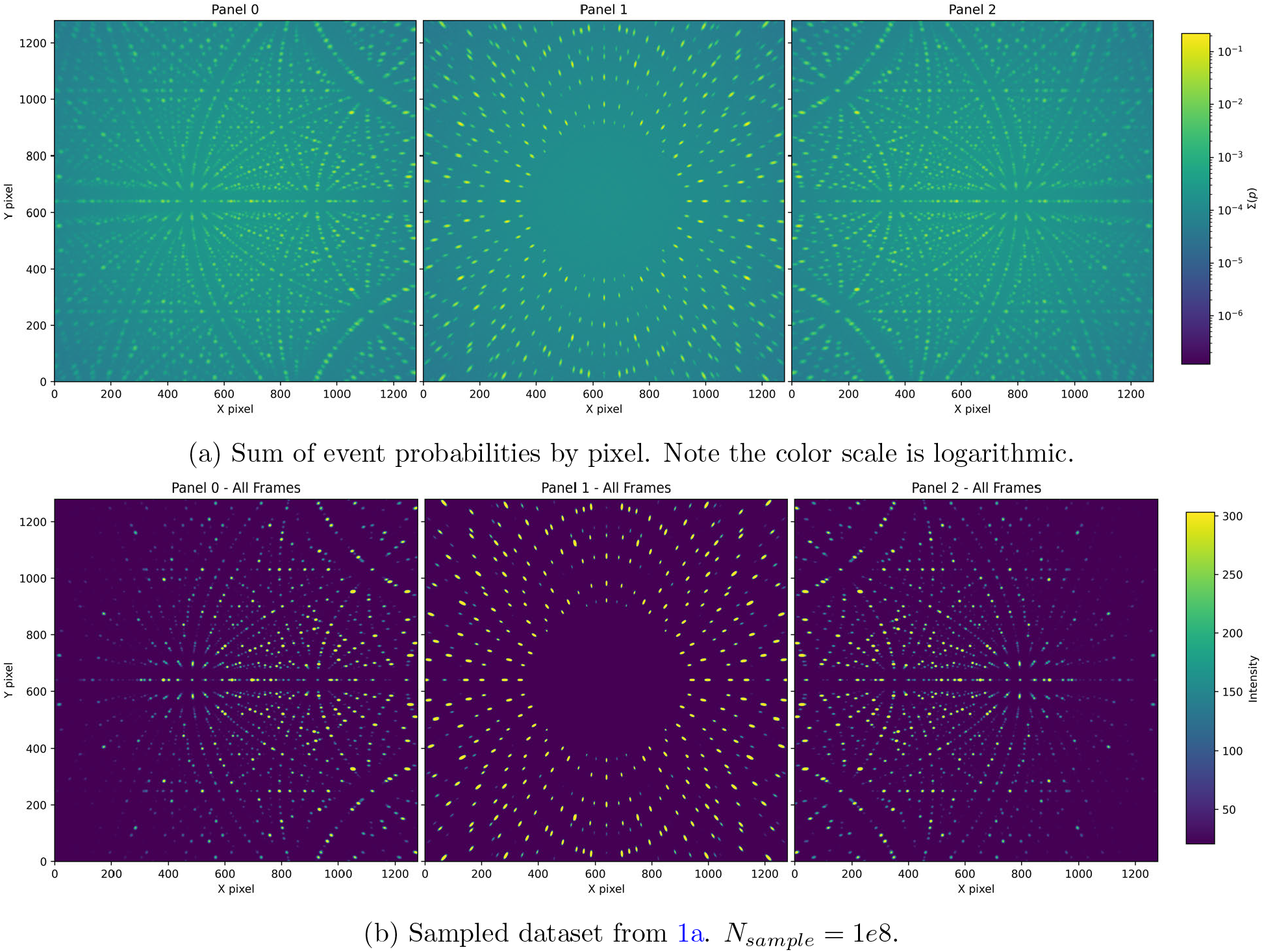
Rubredoxin diffraction simulation with “box” detector geometry. *V*_*crystal*_ = 1*mm*^3^,

The impact of using SPLIT on the Single crystal component is striking (Table 2). SPLIT leads to a proportionate increase in the number of recorded events, with minimal computational overhead. The speed-up of SPLIT comes from caching the list of reflections that are in the Ewald shell (see Section 2.2.1), the calculation of which is expensive with large reflection lists. The SPLIT rays, all of which have the same properties as the original ray and thus would have the same reflections in their Ewald shells, then use the cached list, which is much faster. In addition, using SPLIT avoids re-computation of the neutron ray trajectory from source to sample, further improving efficiency. A SPLIT value of 10000 or more leads to effectively complete sampling of the detector surfaces, as indicated by the jump in min(Σ(*p*)) from 1000 to 10000.

**Table 2:**
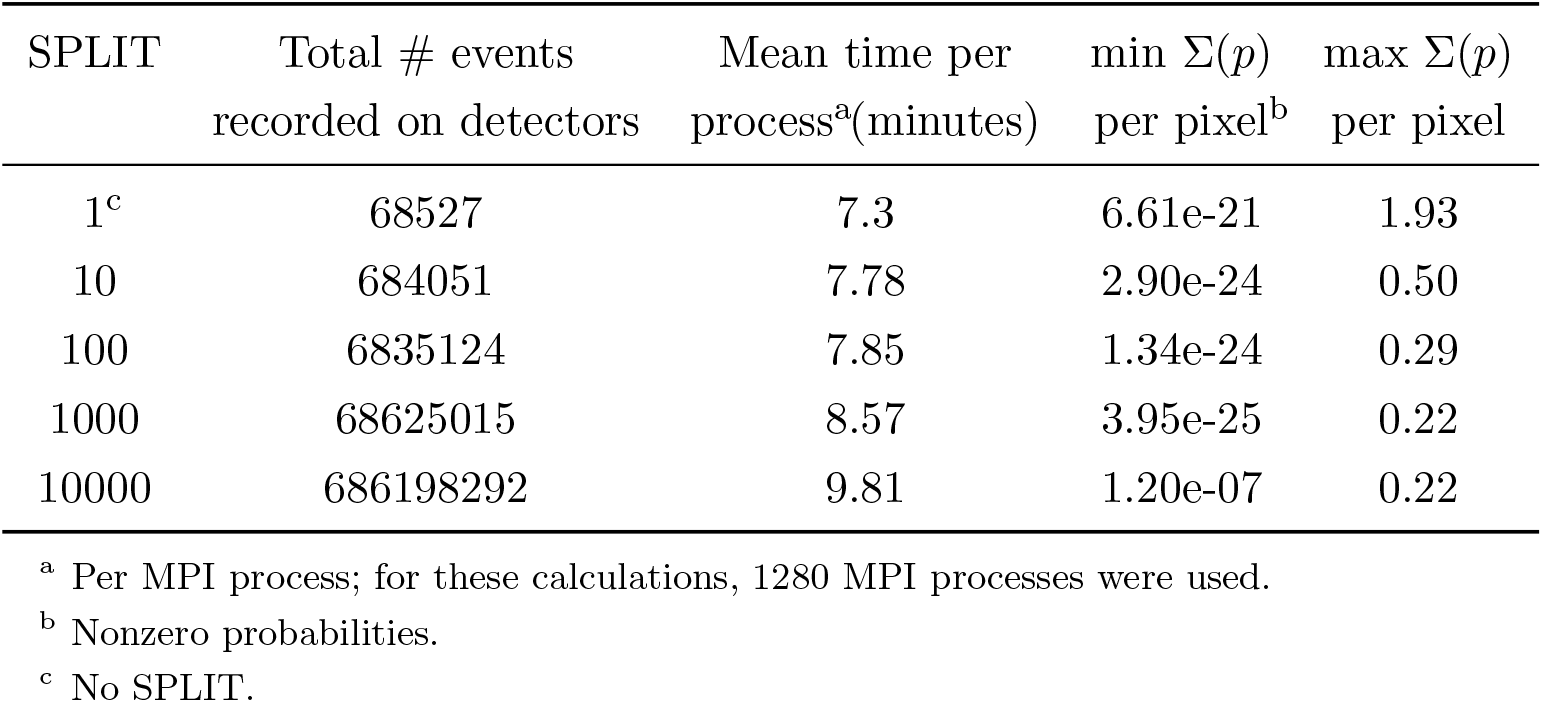
Effects of SPLIT on Simulations of Rubredoxin Diffraction (*N*_*sims*_ = 1*e*11)

A visual depiction of the simulation results certainly looks, to the trained eye, like typical Laue diffraction for a protein crystal. When binned and histogrammed by TOF, each bin looks like monochromatic protein diffraction. A movie of the histogrammed binned by TOF shows the evolution of Bragg diffraction as a function of the incoming neutron ray wavelength (Supporting Information). These encouraging results were confirmed by processing the binned/histogrammed data in DIALS, showing that the simulated dataset is consistent with rubredoxin diffraction. The results are shown in Table 3.

**Table 3:**
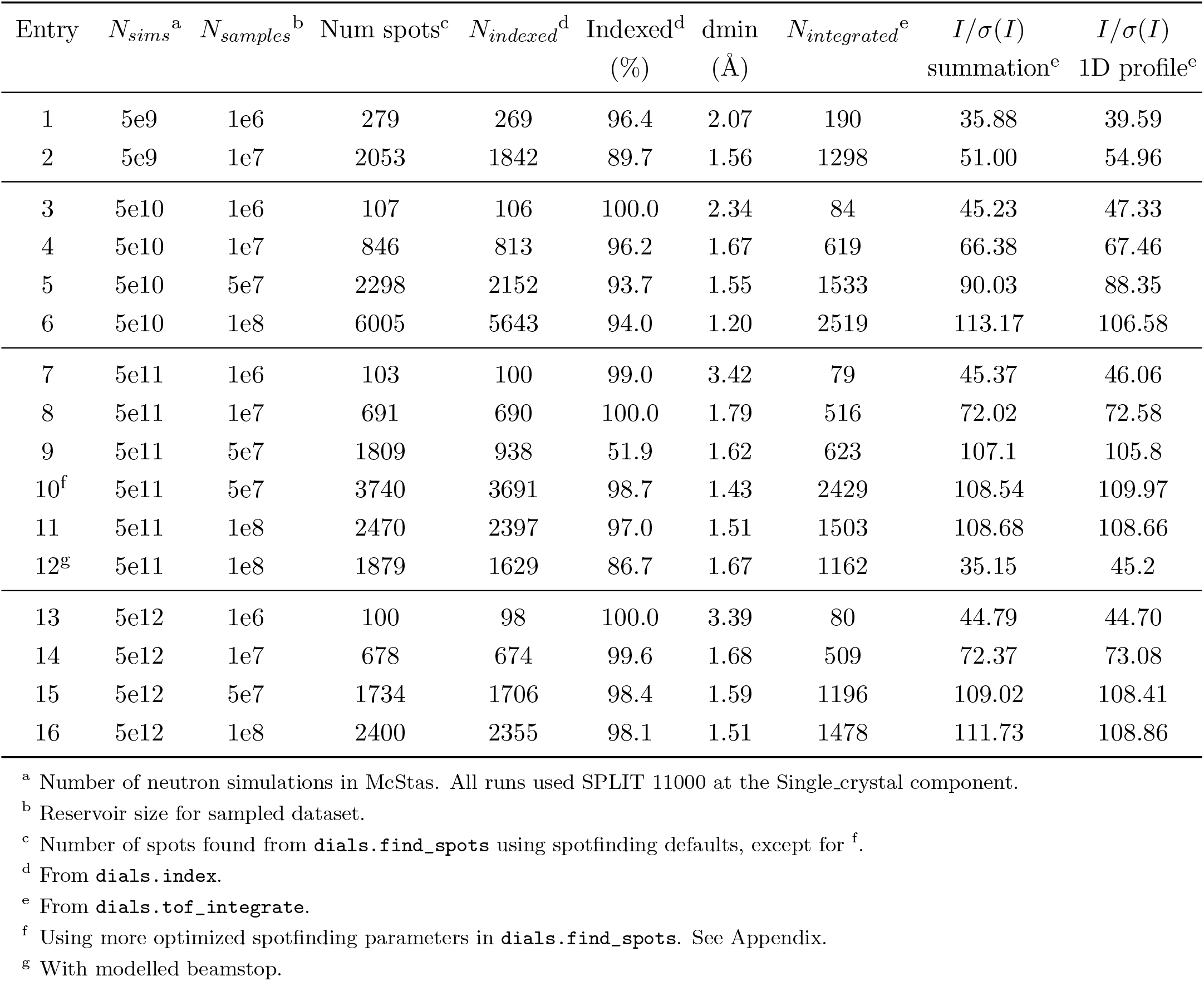
Results of DIALS processing on Simulated Datasets of Rubredoxin Diffraction with Varying *N* _*sims*_ and *N* _*samples*_.

The reservoir sample size *N*_*sims*_ has, unsurprisingly, the most significant effect in data reduction results. Increasing the reservoir sample size is the statistical equivalent of collecting more data, all other things being equal. A reservoir size of *N* = 5*e*7-1*e*8 seems to be the point of diminishing returns. Importantly, sampling is done across events recorded on *all three* detector panels, since the weighted probabilities all derive from the same source (scattering from the source), although McStas separates the results recorded on different monitors. As *N*_*samples*_ increases, so does *d*_*min*_ decrease in turn, in accordance that a larger *N*_*samples*_ corresponds to longer data collection, and weaker reflections take more time to collect. Laue images of simulations of varying sample size can be found in the Supporting information. Assuming that the event list being sampled is randomly distributed (which it is in our case), one can see that the strongest reflections are almost fully sampled even at low *N*, much like what would be expected during data collection.

### 5.2 Simulating Reality: Other sources of scattering

The above, idealistic simulations of Bragg diffraction do not accurately reflect a realistic data collection. Even if the crystal is perfect, other sources of scattering from air, water, the beamstop, inaccurately placed components, etc. contribute to the background of the measurement. Detector jitter, electronic noise, beam anisotropy, *γ*-rays from absorbed neutrons, and cosmic rays are also sources of background in real *n*-MX measurements.

In MLFSOM, Holton et al. attempted to exhaustively model sources of measurement error in X-ray MX experiments, using many semi-empirical relations. *Ab-initio* methods like ours are less amenable to such exhaustive searches, as the computational complexity explodes quickly. This is not to say that sources of noise and background cannot be added after the fact; we will investigate this in the future.

Nonetheless, we can make some inroads on modelling some of the most significant sources of exogenous scattering. We can model a Union beamstop— one containing ^6^LiF in an aluminium tube, 1 cm downstream from the sample— and measure the effects of scattering up to the beamstop component. The results of this simulation are shown in Fig. 2. The effects of extra scattering from air and/or beamstop are particularly prominent around *d* ≈ 1.5Å, as indicated by the diffuse scattering cone there. Issues with sampling can also arise, as transmitted beam, or low-angle scattering, near the beamstop (centre of Panel 1) leads to extremely high neutron probabilities. A “mask” around the beam centre, should a detector panel pass through it, or a higher sampling reservoir size, can overcome the high probabilities of the few affected pixels.

**Figure 2.**
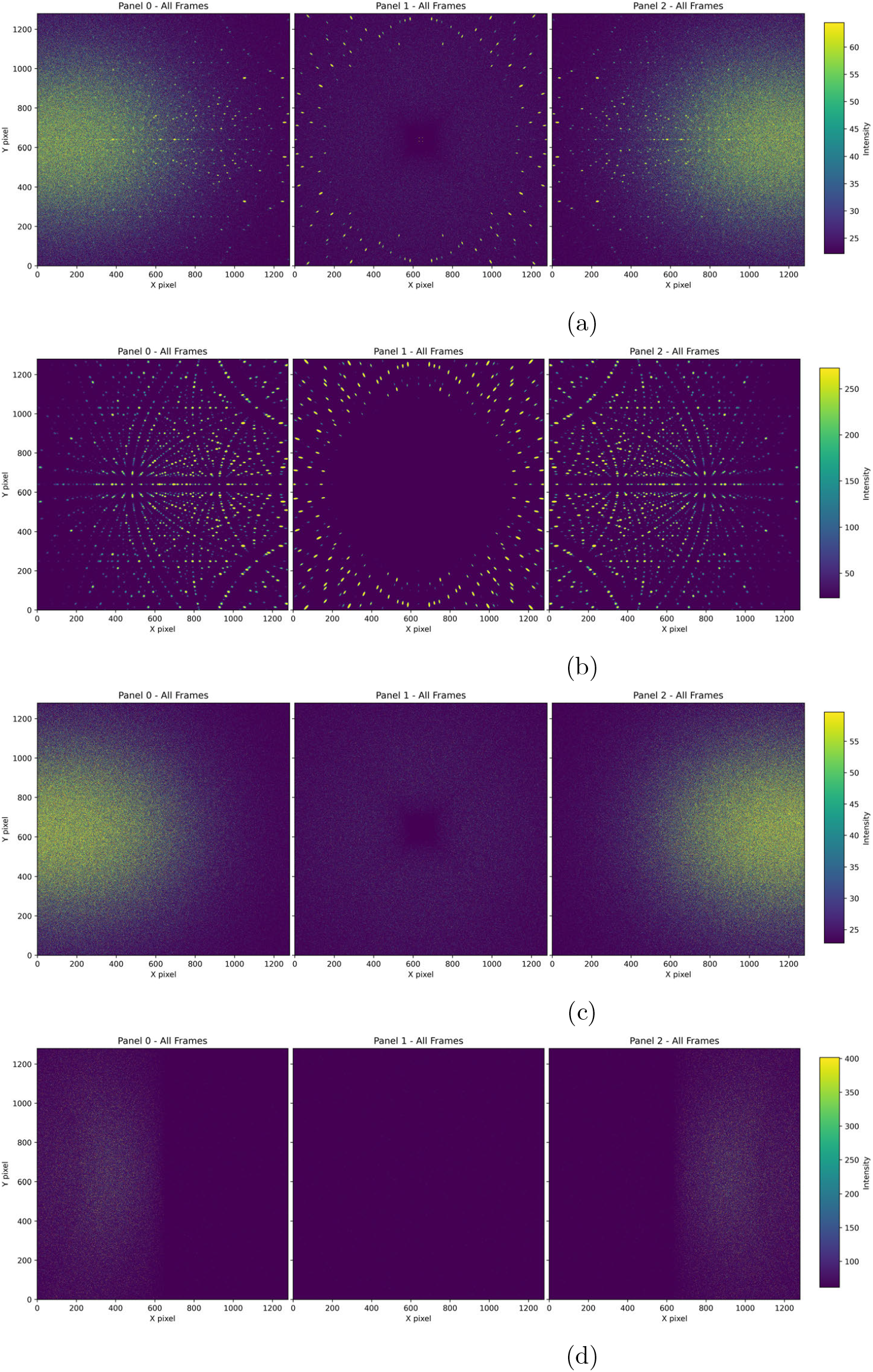
Rubredoxin simulation with “box” detector geometry, with a 0.35 mm ^6^LiF beamstop in a thin Al tube placed 1 cm downstream of the sample. *V*_*crystal*_ = 1*mm*^3^, *N*_*simulations*_ = 5*e*11, *SPLIT* = 11000, *N*_*sampled*_ = 1*e*8. Sampling from four different conditional monitors: (a) monitoring all events, regardless of source; (b) monitoring scattering from crystal; (c) monitoring scattering from air; (d) monitoring scattering from beamstop.

Modification of the SPLIT method for the Union beamstop simulation was necessary to avoid erroneous random events of high probability— on average 1000 times higher than other scattering events. Instead of setting a SPLIT on the Single crystal component alone, the SPLIT value was divided between the Union Master and the Single crystal components, such that the product of the two SPLIT parameters was 11000 (e.g. setting SPLIT 11 for the Union component and 1000 for the Single crystal). This gave more reasonable results.

We can distinguish between events created from crystal scattering and elsewhere by using conditional monitors in McStas that only record events when certain conditions are met, e.g. a neutron scattering off the crystal, or air, or both (or neither). Fig 2 shows the results of an NMX simulation with rubredoxin crystal and a Union beamstop using conditional monitors: one from scattering off the crystal (Fig. 2b), air (Fig. 2c), and the beamstop (Fig. 2d). Clearly, air is the biggest contributor to exogenous scatter, while the beamstop’s contribution is relatively small. Conditional monitors will be invaluable when simulating more complex scenarios, such as the presence of H_2_O.

### 5.3 Simulating the Detector Geometries of NMX

So far, the simulations discussed herein have been in a simple “box” geometry, with the three detector panels huddled around the sample, about 35 cm away (see Supporting Information). We have simulated rubredoxin diffraction with all 11 detector configurations, and processed the resulting TOF-Laue data with DIALS; the detailed results are in the Supporting Information. It is possible to model all 33 detector positions at once due to the fact that a monitor in McStas simply records any neutrons that pass through it, and can do so unimpeded if desired. The ability to do this will prove very useful for data collection strategy and planning. Notably, one can see that, as expected, the effects of air scatter become significantly more severe for detector geometries where the panels are very far from the sample. Even with the same *N*_*samples*_, in configurations with detectors furthest away from the sample, like configurations 9 and 10, exogenous scatter dominates the sampled data.

## 6 Conclusions

We have described our initial work on simulations of the TOF-Laue *n*-MX experiment on the NMX Macromolecular Diffractometer at ESS, using McStas. With the new single-pass weighted random sampling with reservoir, we can convert the McStas event probability output into realistic data, as indicated by processing these data with DIALS. We will use these methods as a template for instrument commissioning, as well as data collection strategies and analysis.

## A Typical Simulation Workflow

1.Generate the relevant .lau file for Bragg reflections, using the calculate structure factors.ipynb file. Save the .lau file in the same folder as the McStas `NMX.instr` file for the McStas simulation.

2.Run `mcrun`, an automated process that generates the McStas C-code, runs the simulation binary, and handles the output. This step is usually done on HPC systems:

~~~
mcrun -c --mpi=auto NMX.instr --format=NEXUS \
--IDF -n{Nsims} XtalPhiY=0 -d /path/to/output
~~~

The `-c` option forces recompilation of the binary. The `--IDF` instruction outputs the Mantid-style XML data for the source and monitor components. Setting a McStas variable declared in `NMX.instr`, such as `XtalPhiY=0`, on the command line ensures that all defaults are chosen without querying the user (necessary for batch processing). The output directory, set by `-d`, must not already exist or `mcrun` will exit. The NeXus output is saved in

`/path/to/output/mccode.h5`.

3. Sample the McStas data:

~~~
python /path/to/stream_sampling_mcstas/sampling_to_nexus.py \
--do-histogram --n-samples 100_000_000 -i path/to/output/mccode.h5 -o sampling_out
~~~

This will generate a NeXus file, `sampling_output.h5`, in TOFRaw format.

4. Bin and histogram the sampled data. Usually 50 bins is sufficient.

~~~
essnmx-reduce --input-file sampling_output.h5 \
--output-file scipp_output.h5 --nbins 50
~~~

5. Process the binned/histogrammed data with DIALS:

~~~
dials.import scipp_output.h5
dials.find_spots imported.expt
dials.index imported.expt strong.refl \
   unit_cell=“34.37,35.18,43.99,90,90,90” space_group=19
dials.refine indexed.expt indexed.refl
dials.tof_integrate refined.expt refined.refl method=profile1d \
   profile1d.n_restarts=500 profile1d.min_beta=1e-5 profile1d.min_alpha=1e-5 \
   profile1d.max_beta=10 profile1d.max_alpha=10
~~~

We include the unit cell and symmetry for indexing, since they are known, but it is usually not necessary. In cases where there is little noise in the data, such as in the case of the “ideal” beamstop, running `dials.find_spots` with default parameters sometimes gives erroneous results. Setting the following parameters can help:

~~~
spotfinder {
  threshold
    { dispersion {
      gain = 0.01
      kernel_size = 50 50
      sigma_background = 1
      sigma_strong = 2
      min_local = 1
      global_threshold = 0
}
      algorithm=radial_profile
}
filter {
    min_spot_size = 30
    max_spot_size = 9000
    max_separation = 20
}
}
~~~

## Supporting information

Calculating Structure factors in Jupyter

McStas instrument file for NMX

McStas instrument file for NMX

## Acknowledgements

The authors would like to thank Daniel Lundström for providing the detector configuration pictures; Massimiliano Novelli for assistance with data archival; and Zoë Fisher, Swati Aggarwal, and Dorothea Pfeiffer for helpful discussions.

## Data availability

Primary McStas results from simulations can be accessed via SciCat through the following DOI: 10.17199/d9ca04b5-84e8-4428-ac75-ebf41066dfc7

A Jupyter notebook, calculate structure factors.ipynb, is available in the Supporting Information. This notebook demonstrates how to generate the list of structure factors in Crystallographica .lau format from PDB entries.

Two McStas instrument files for NMX-one with a simple “box” geometry and ideal beamstop, and one with all detector configurations and a Union beamstop, are available in the Supporting Information.

The scipp/essnmx software can be found at https://doi.org/10.5281/zenodo.10731810

The StreamSampling.jl software can be found at https://doi.org/10.5281/zenodo.12826684

The stream sampling mcstas software can be found at https://doi.org/10.5281/zenodo.19137556

## Footnotes

1 Currently we expect to be open to users in 2027.

2 We discard the energy of the neutron as we do not measure it in our detectors— only the TOA and position on the detector. The energy is instead calculated from the TOA.

3 Nota bene about memory: more complicated computations require more memory. The major source of memory consumption for our computations is the reflection list for protein crystals. Each MPI process loads the entire reflection list into memory, so for large or very high-resolution protein crystals, the amount of memory required skyrockets into unreasonable territory quickly, and out-of-memory errors can occur. We are currently working on improving this in McStas, but it is currently *only* an issue when doing massively parallel computations on HPC resources.

4 Compressing large lists of floats of relatively uncorrelated data, such as what McStas outputs, is inefficient.

