## Supplemental Information for "Simulating Neutron Protein Crystallography Experiments: Applications to the Development of the NMX Instrument at ESS"

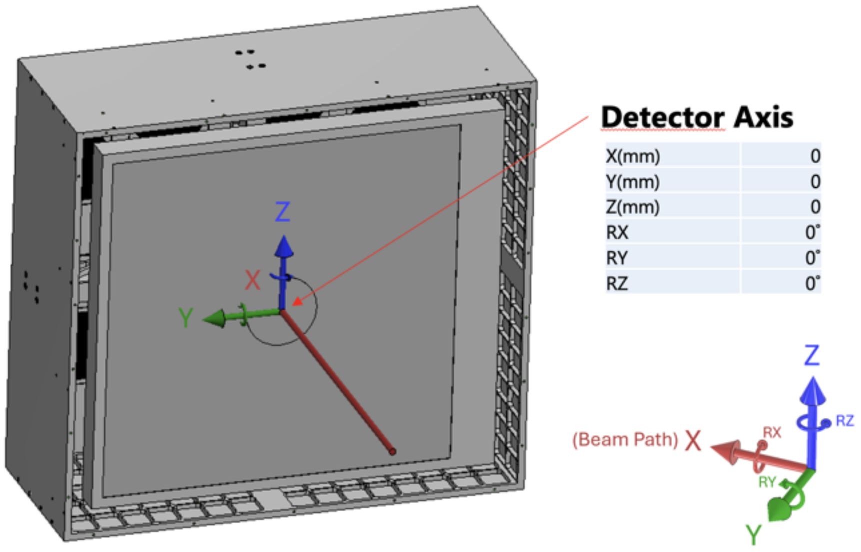


Figure S1: Coordinate system for detector panels. The X-axis is the neutron beam path. The Z-axis is parallel to gravity. The origin of the axis labels corresponds to the sample position. Measurements in Tables S1 and S2 are relative to the center of the detector panel. Rotations are according to the right-hand rule.

Table S1: Positions of Detectors for NMX Configurations (Box and Configurations 1–5)

| Panel | | Box | C1 | C2 | C3 | C4 | C5 |
|---|---|---|---|---|---|---|---|
| Panel 1 | X (mm) | 250 | 270 | 331 | 336 | 581 | 451 |
| | Y (mm) | 0 | 0 | 0 | 0 | 0 | 0 |
| | Z (mm) | 0 | 0 | 160 | 0 | 0 | 0 |
| | RX (°) | -90 | 0 | 0 | 135 | 0 | 0 |
| | RY (°) | 0 | 0 | 0 | 0 | 0 | 0 |
| | RZ (°) | 0 | 0 | 0 | 0 | 0 | 0 |
| Panel 2 | X (mm) | 320 | 85.85 | 0 | 0 | 241.61 | 187.68 |
| | Y (mm) | 0 | 235.86 | 331 | 237.59 | 470.25 | 0 |
| | Z (mm) | 0 | 160 | 160 | 237.59 | 0 | 464.52 |
| | RX (°) | 0 | 180 | 180 | 180 | 180 | 0 |
| | RY (°) | 0 | 0 | 0 | -45 | 0 | -68 |
| | RZ (°) | 0 | 110 | 90 | 90 | 58 | 0 |
| Panel 3 | X (mm) | 250 | -113 | 0 | 0 | 241.61 | -318.91 |
| | Y (mm) | 0 | -386.49 | -501 | -237.59 | -470.25 | 0 |
| | Z (mm) | 0 | 122.85 | 160 | 237.59 | 0 | 318.91 |
| | RX (°) | 90 | 0 | 0 | 180 | 0 | 0 |
| | RY (°) | 0 | -17 | 0 | -45 | 0 | -135 |

| Panel | | Box | C1 | C2 | C3 | C4 | C5 |
|---|---|---|---|---|---|---|---|
| | RZ (°) | 0 | -110 | -90 | -90 | -58 | 0 |

Table S2: Positions of Detectors for NMX Configurations (Configurations 6–11)

| Panel | | C6 | C7 | C8 | C9 | C10 | C11 |
|---|---|---|---|---|---|---|---|
| Panel 1 | X (mm) | 435.63 | 1001 | 848.9 | 1001 | 1001 | 342.36 |
| | Y (mm) | 116.73 | 0 | -530.45 | 0 | 0 | 0 |
| | Z (mm) | 0 | 0 | 0 | 0 | 0 | 940.63 |
| | RX (°) | 180 | 0 | 0 | 0 | 0 | 0 |
| | RY (°) | 0 | 0 | 0 | 0 | 0 | -70 |
| | RZ (°) | 15 | 0 | -32 | 0 | 0 | 0 |
| Panel 2 | X (mm) | 319.57 | 799.43 | 848.9 | -707.81 | 799.43 | -309.33 |
| | Y (mm) | -424.08 | 907.21 | 530.45 | 707.81 | 0 | 0 |
| | Z (mm) | 0 | 0 | 0 | 0 | 602.42 | 952.01 |
| | RX (°) | 0 | 0 | 0 | 0 | 0 | 0 |
| | RY (°) | 0 | 0 | 0 | 0 | 37 | -108 |
| | RZ (°) | -53 | 65 | 32 | 135 | 0 | 0 |
| Panel 3 | X (mm) | -240.5 | -707.81 | -121.99 | 423.04 | 259.08 | -819.97 |
| | Y (mm) | -416.56 | -707.81 | -993.54 | -907.21 | 0 | 0 |
| | Z (mm) | 0 | 0 | 0 | 0 | 966.89 | 574.15 |
| | RX (°) | 0 | 0 | 0 | 0 | 0 | 0 |
| | RY (°) | 0 | 0 | 0 | 0 | 75 | -145 |
| | RZ (°) | -120 | -135 | -97 | -65 | 0 | 0 |

The following Python code can be used to convert the above measurements (in ESS coordinate system) to McStas coordinates:

```
def coords_to_mcstas(x,y,z,rotx,roty,rotz):
    from scipy.spatial.transform import Rotation as R
    x2, y2, z2 = y, z, x  # xyz (ESS) = YZX (McStas)
    rotation = R.from_euler("xyz", [rotx, roty, rotz], degrees=True)
    ang = rotation.as_euler("YZX", degrees=True)
    return x2, y2, z2, ang[0], ang[1], ang[2]
```

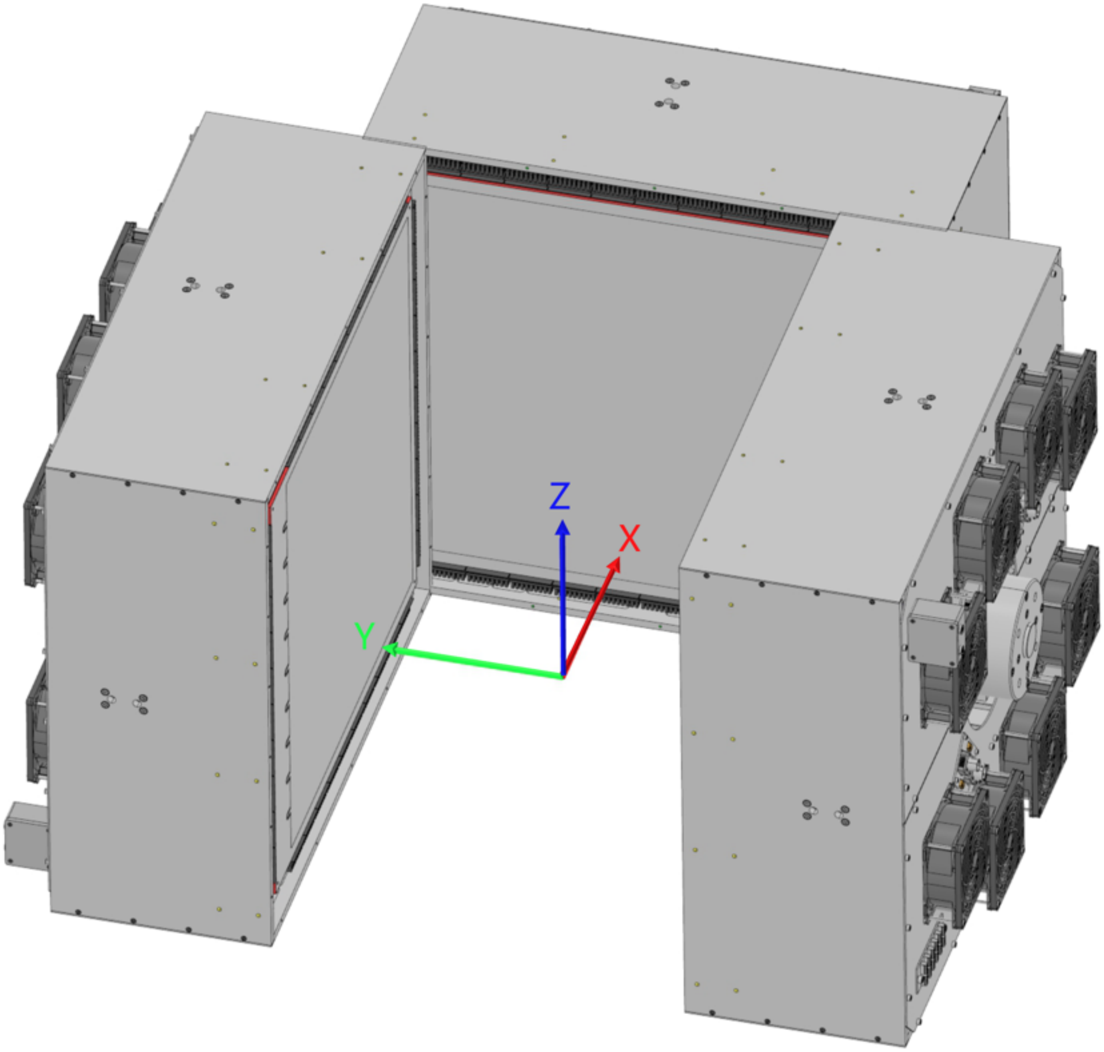


Figure S2: Detector "Box" Configuration.

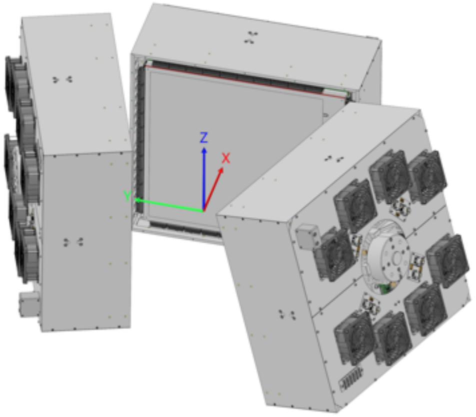


(a)

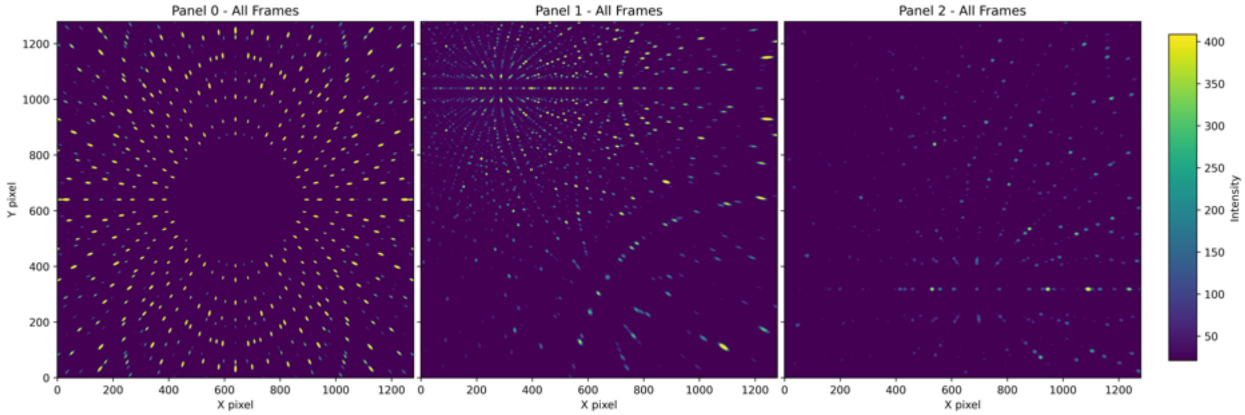


(b)

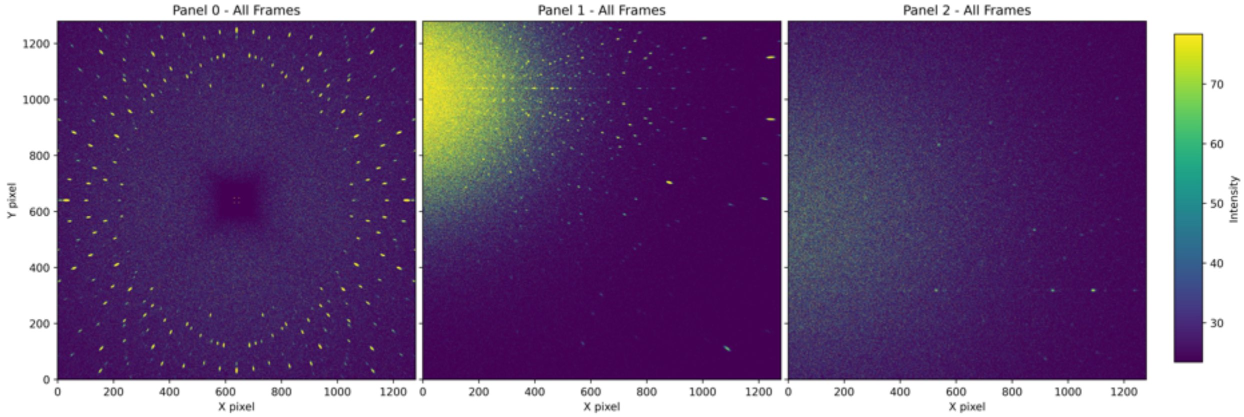


(c)

Figure S3: (a) Configuration 1. The X-axis is the beam path. The sample position is the origin of the axis labels. (b) Simulation of crystal only. (c) Simulation with beamstop. $N_{simulations} = 1e11$, $N_{samples} = 1e8$.

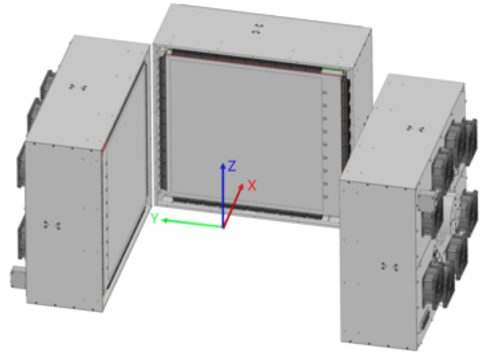


(a)

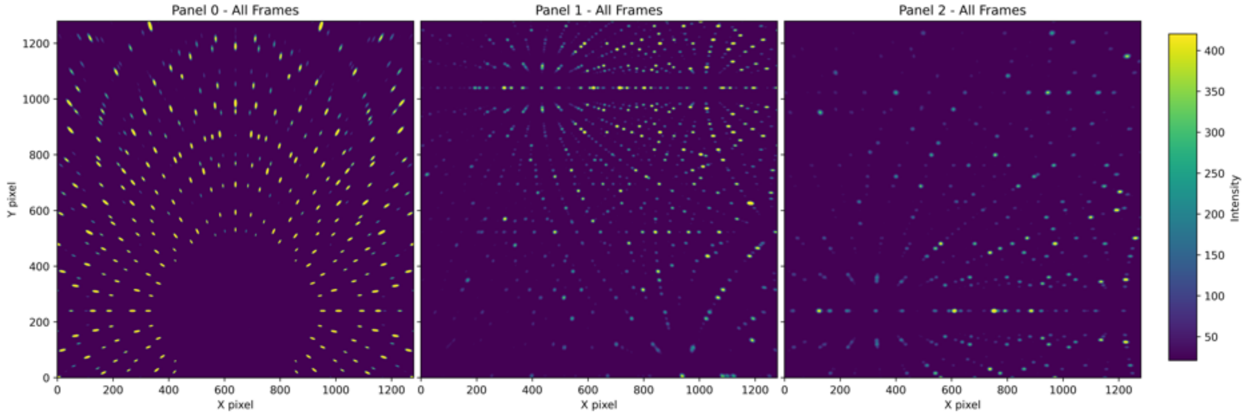


(b)

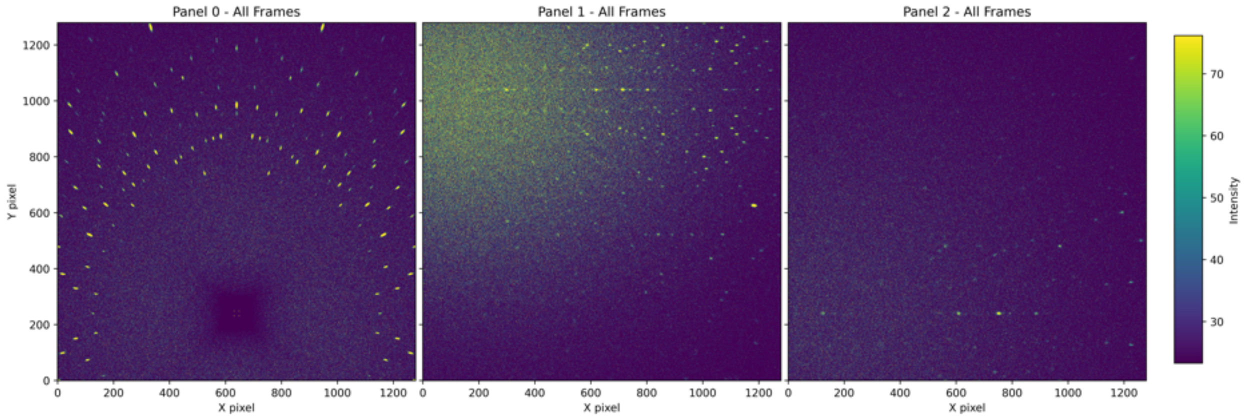


(c)

Figure S4: (a) Configuration 2. The X-axis is the beam path. The sample position is the origin of the axis labels. (b) Simulation of crystal only. (c) Simulation with beamstop. $N_{simulations} = 1e11$, $N_{samples} = 1e8$.

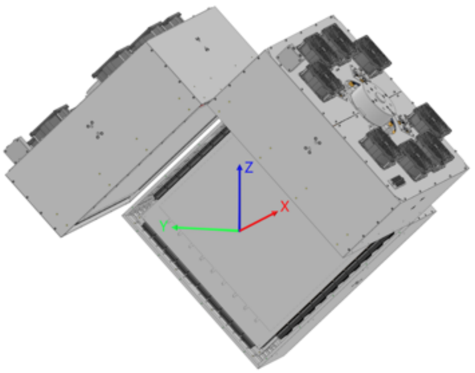


(a)

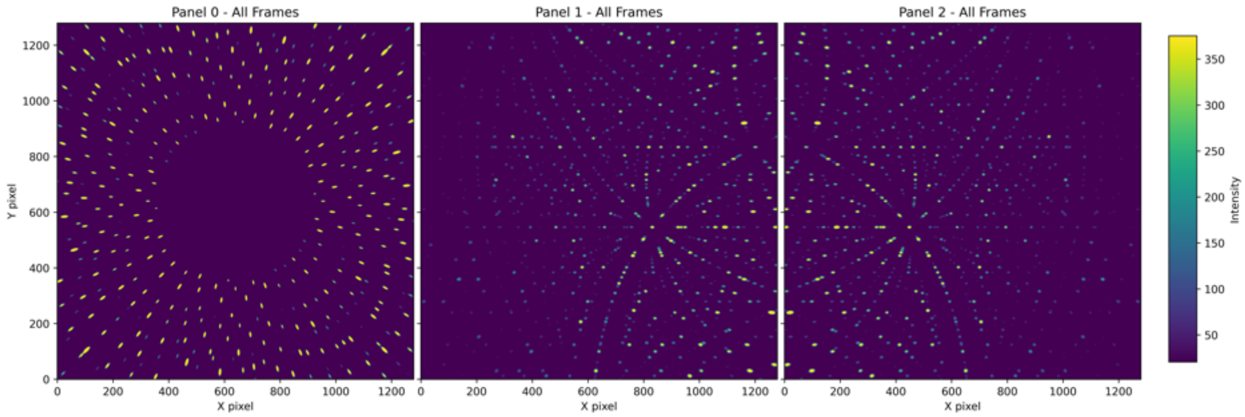


(b)

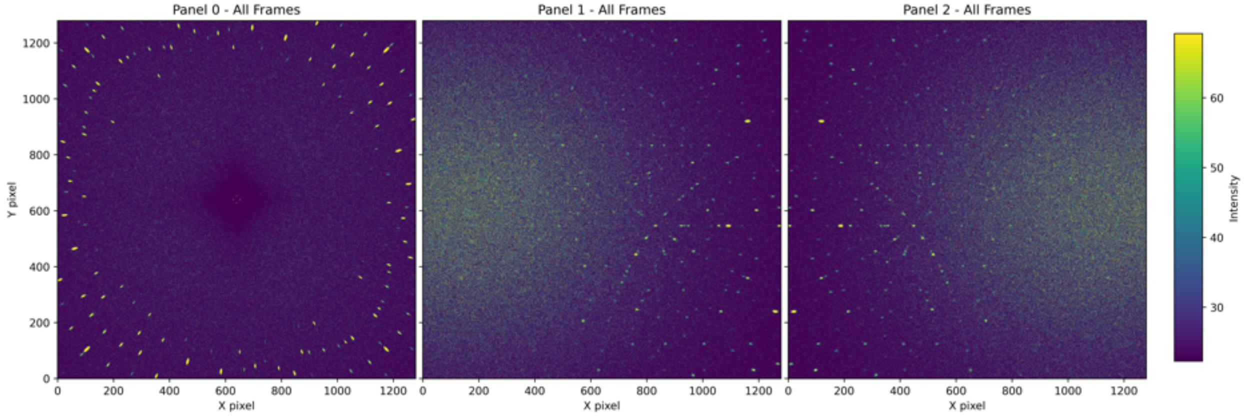


(c)

Figure S5: (a) Configuration 3. The X-axis is the beam path. The sample position is the origin of the axis labels. (b) Simulation of crystal only. (c) Simulation with beamstop. $N_{simulations} = 1e11$, $N_{samples} = 1e8$.

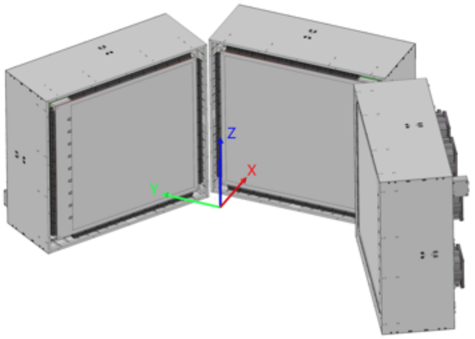


(a)

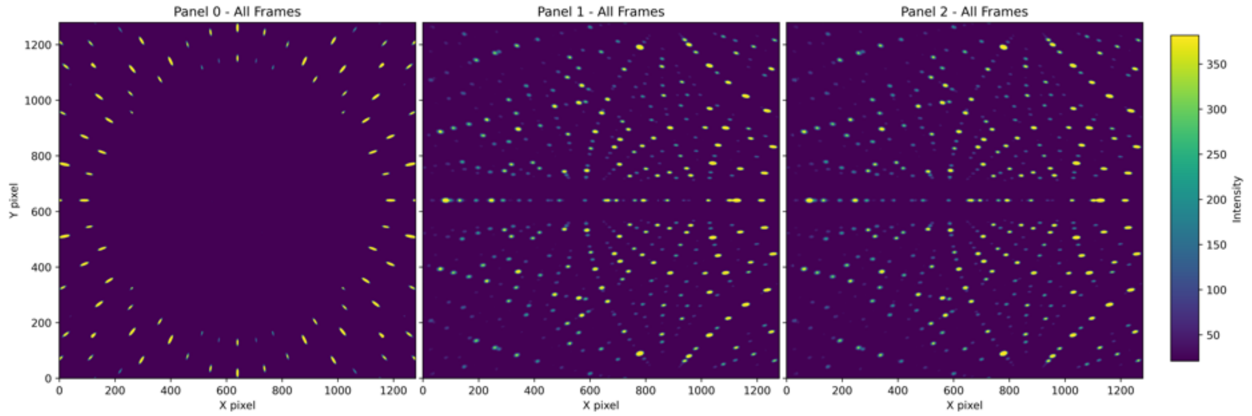


(b)

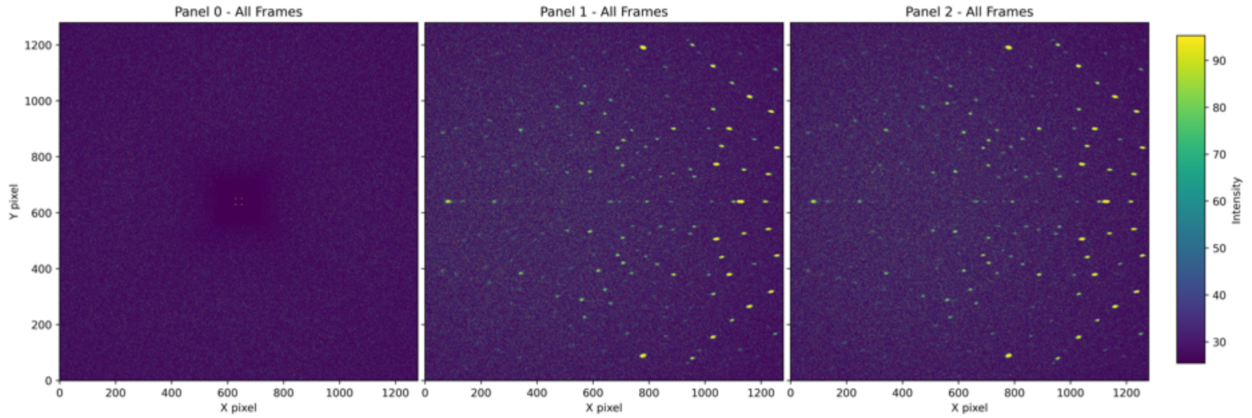


(c)

Figure S6: (a) Configuration 4. The X-axis is the beam path. The sample position is the origin of the axis labels. (b) Simulation of crystal only. (c) Simulation with beamstop. $N_{simulations} = 1e11$, $N_{samples} = 1e8$.

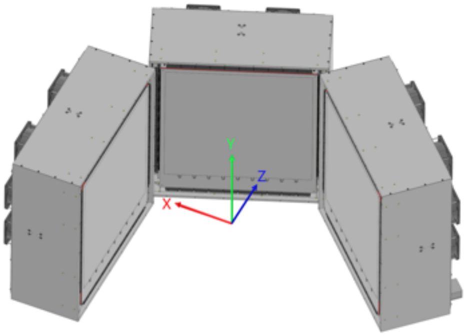


(a)

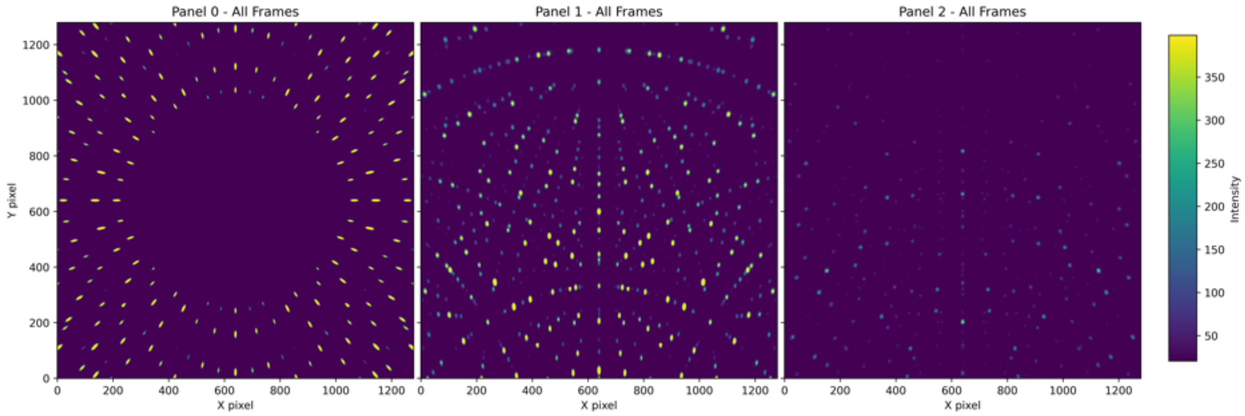


(b)

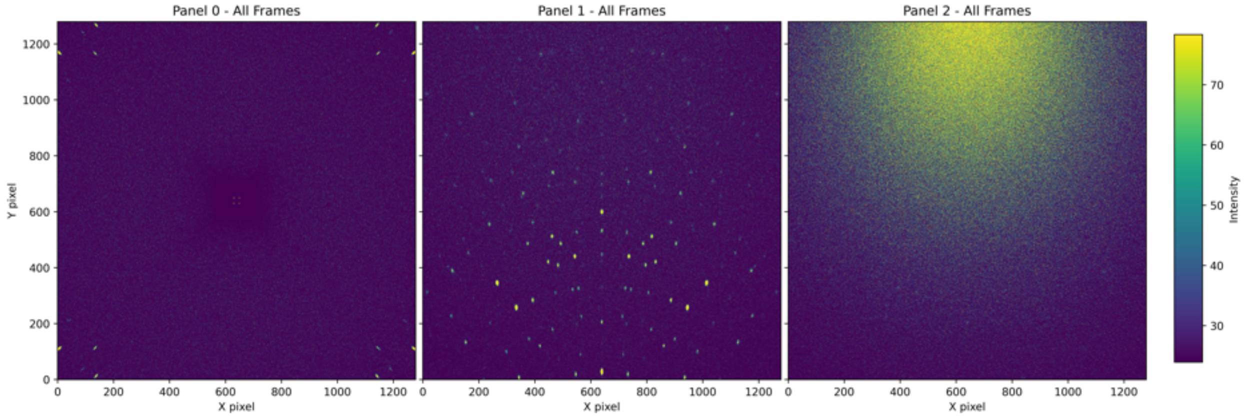


(c)

Figure S7: (a) Configuration 5. The X-axis is the beam path. The sample position is the origin of the axis labels. (b) Simulation of crystal only. (c) Simulation with beamstop. $N_{simulations} = 1e11$, $N_{samples} = 1e8$.

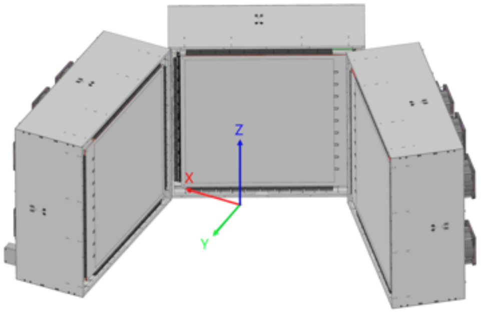


(a)

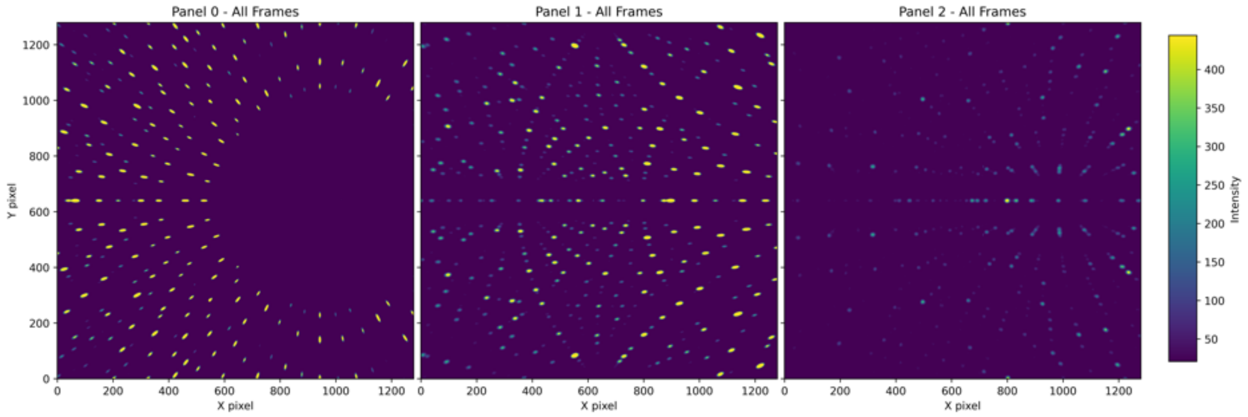


(b)

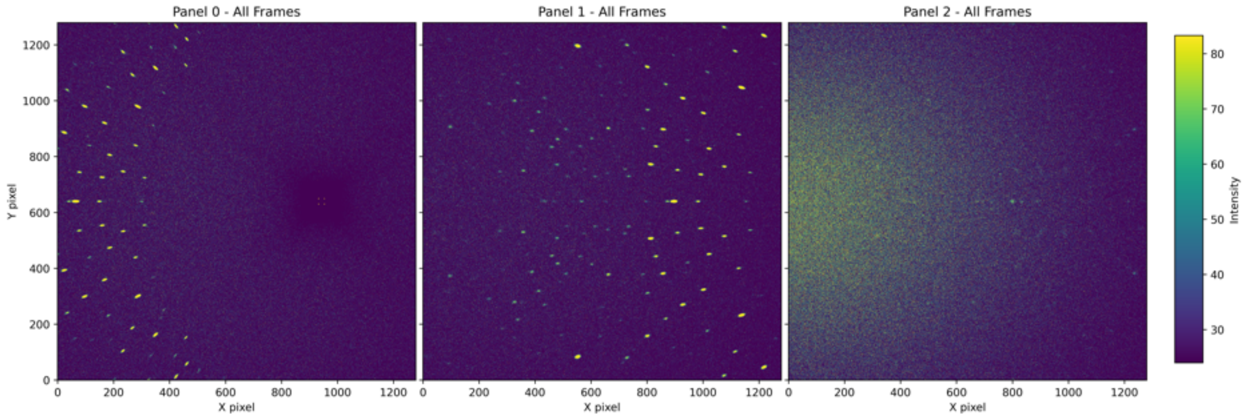


(c)

Figure S8: (a) Configuration 6. The X-axis is the beam path. The sample position is the origin of the axis labels. (b) Simulation of crystal only. (c) Simulation with beamstop. $N_{simulations} = 1e11$, $N_{samples} = 1e8$.

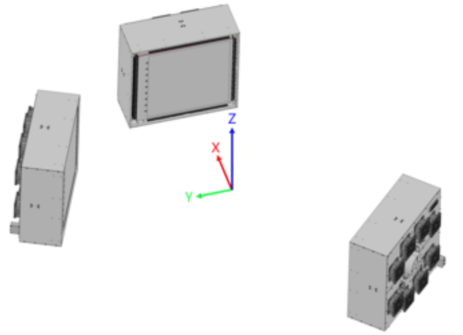


(a)

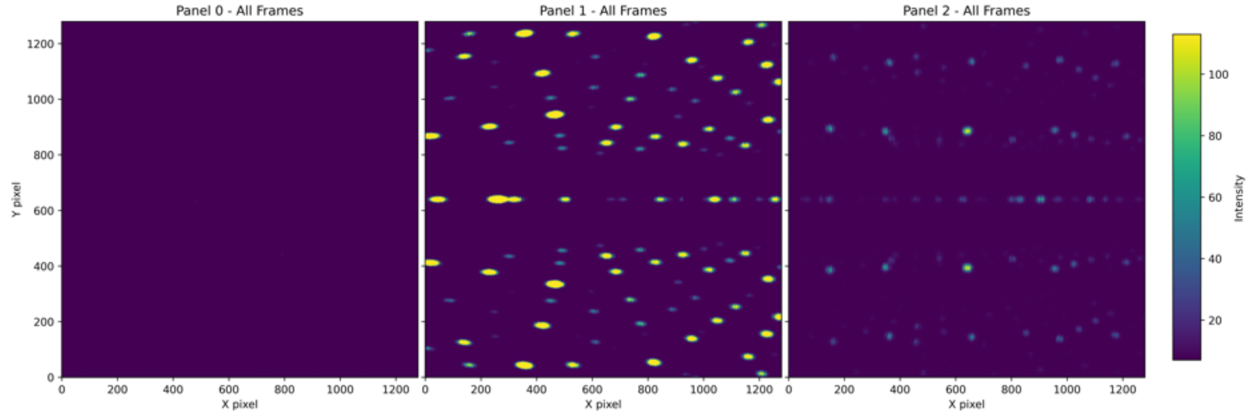


(b)

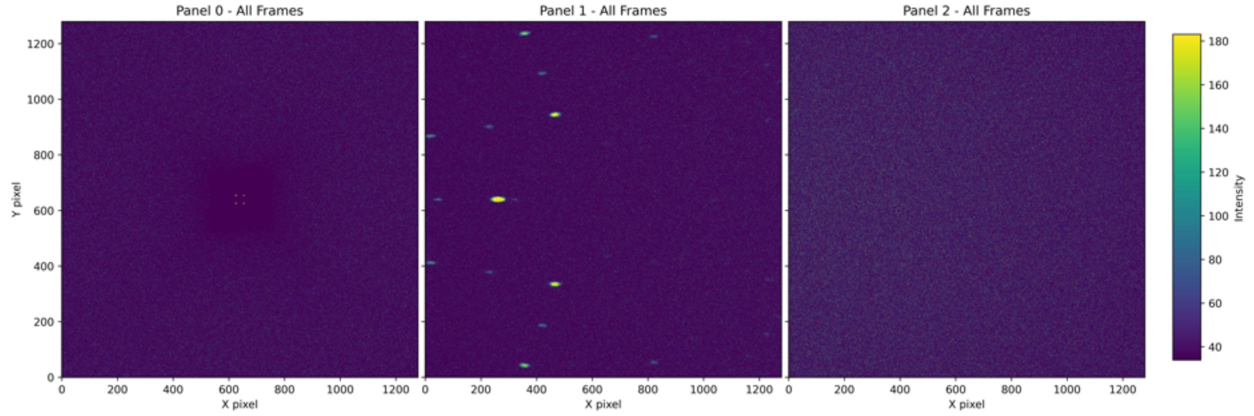


(c)

Figure S9: (a) Configuration 7. The X-axis is the beam path. The sample position is the origin of the axis labels. (b) Simulation of crystal only. (c) Simulation with beamstop. $N_{simulations} = 1e11$, $N_{samples} = 1e8$.

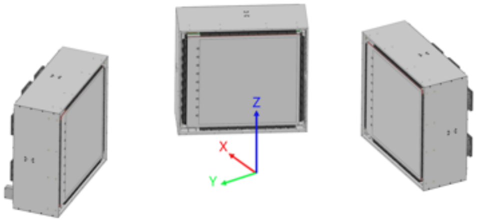


(a)

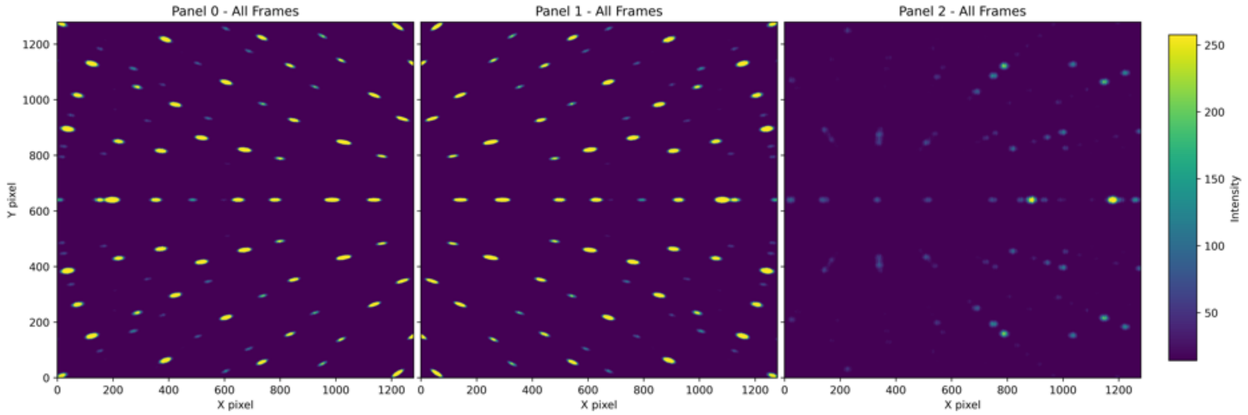


(b)

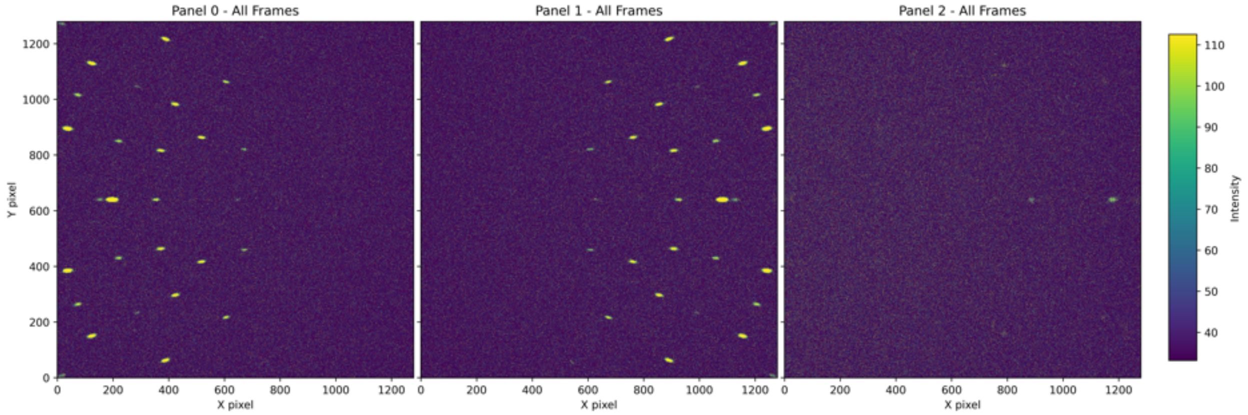


(c)

Figure S10: (a) Configuration 8. The X-axis is the beam path. The sample position is the origin of the axis labels. (b) Simulation of crystal only. (c) Simulation with beamstop. $N_{simulations} = 1e11$, $N_{samples} = 1e8$.

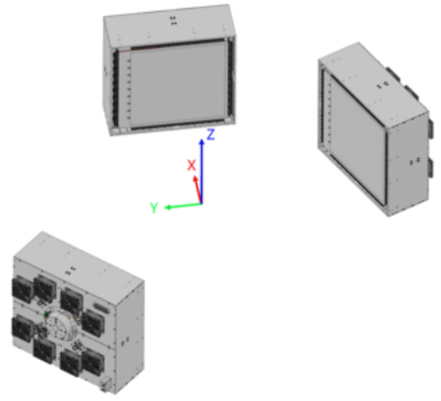


(a)

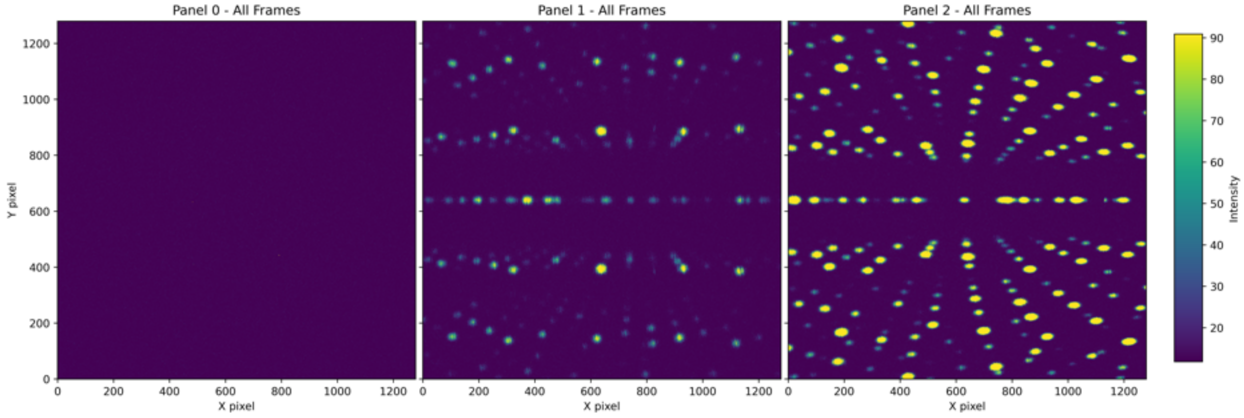


(b)

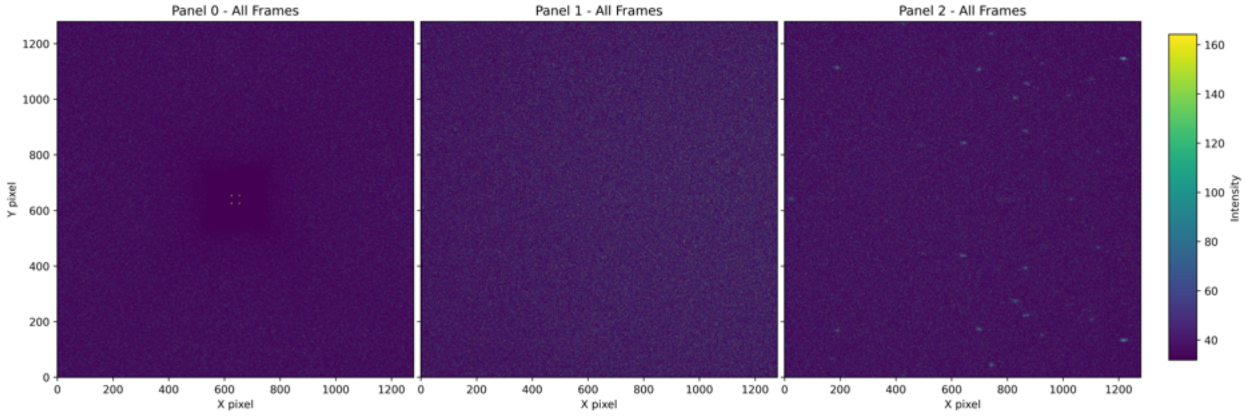


(c)

Figure S11: (a) Configuration 9. The X-axis is the beam path. The sample position is the origin of the axis labels. (b) Simulation of crystal only. (c) Simulation with beamstop. $N_{simulations} = 1e11$, $N_{samples} = 1e8$.

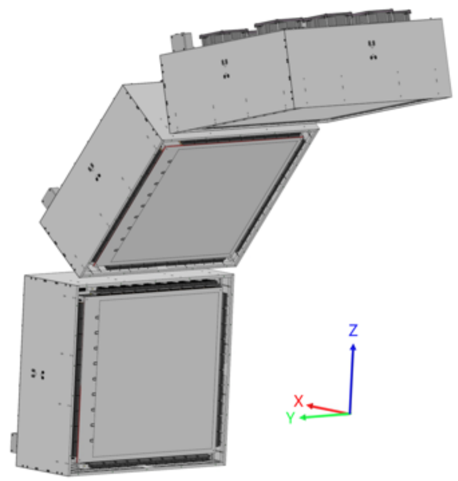


(a)

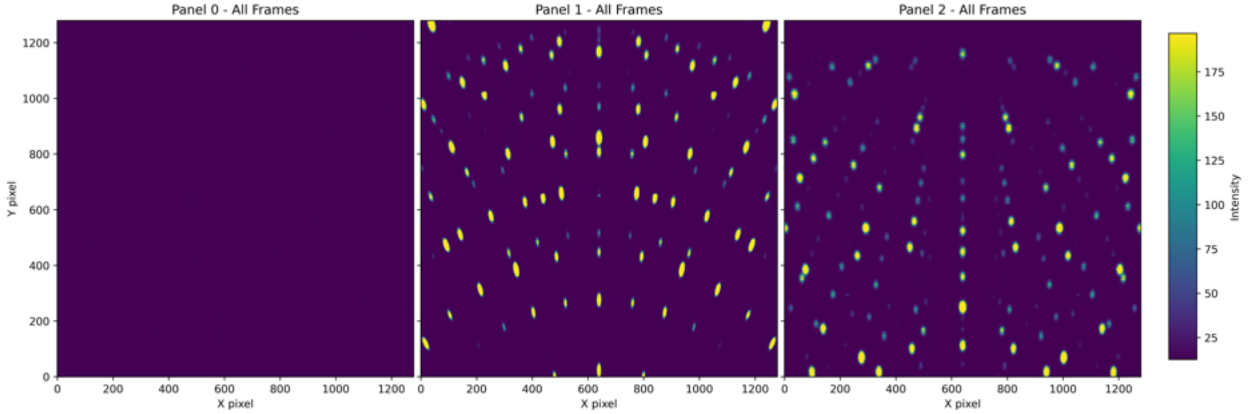


(b)

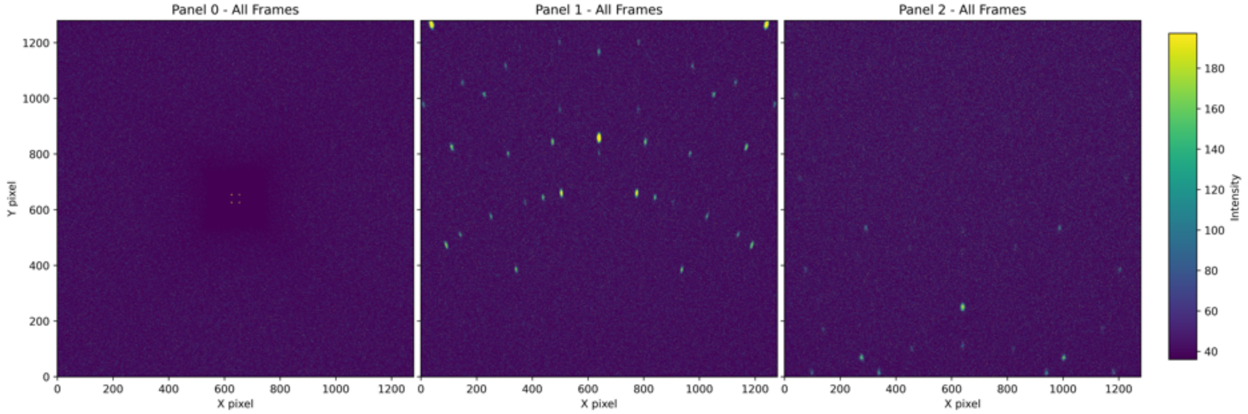


(c)

Figure S12: (a) Configuration 10. The X-axis is the beam path. The sample position is the origin of the axis labels. (b) Simulation of crystal only. (c) Simulation with beamstop. $N_{simulations} = 1e11$, $N_{samples} = 1e8$.

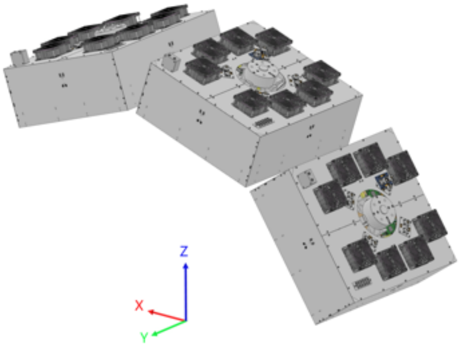


(a)

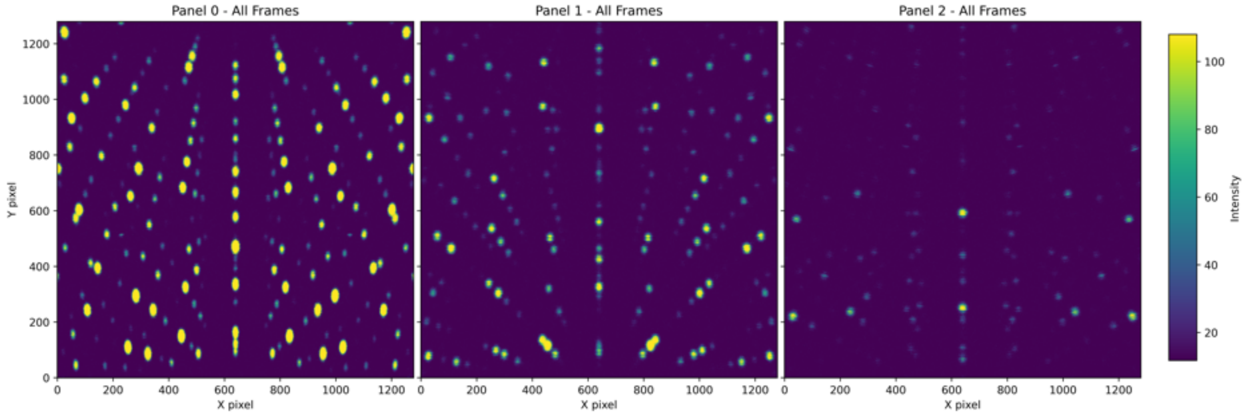


(b)

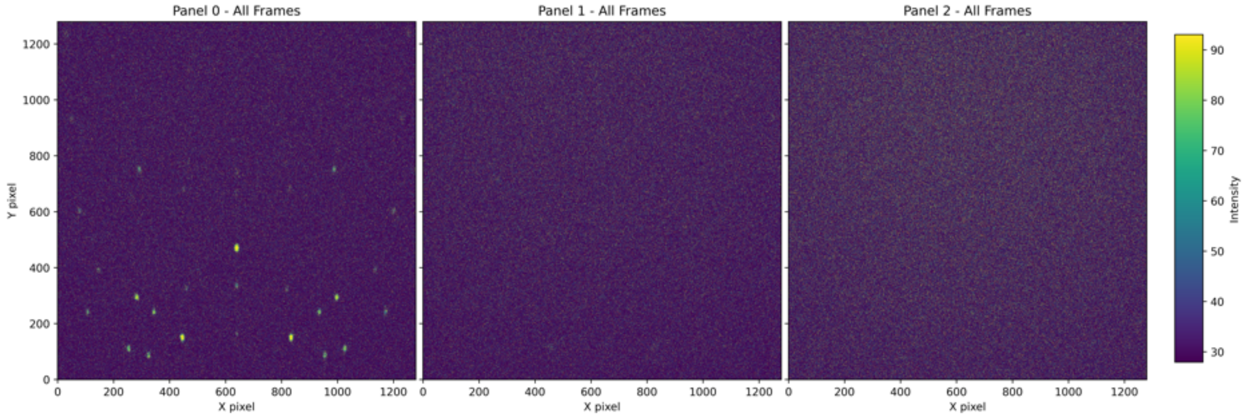


(c)

Figure S13: (a) Configuration 11. The X-axis is the beam path. The sample position is the origin of the axis labels. (b) Simulation of crystal only. (c) Simulation with beamstop. $N_{simulations} = 1e11$, $N_{samples} = 1e8$.

Table S3: Results of DIALS processing of Rubredoxin simulation of NMX detector configurations[a]

| Entry | Configuration | Num spots[b] | $N_{indexed}$[c] | Indexed %[c] | dmin (Å) | $N_{int}/I/\sigma(I)$ summation[d] | $N_{int}$[d]$/I/\sigma(I)$ profile fitting[d] |
|---|---|---|---|---|---|---|---|
| 1 | 1 | 2258 | 2077 | 92.0 | 1.42 | 1291/125.2 | 1280/144.9 |
| 2[e] | 1 | 1241 | 1127 | 90.8 | 1.59 | 839/36.15 | 708/41.49 |
| 3 | 2 | 1531 | 1309 | 85.5 | 1.59 | 810/151.38 | 805/176.06 |
| 4[e] | 2 | 801 | 724 | 90.4 | 1.67 | 592/38.17 | 549/42.44 |
| 5 | 3 | 1735 | 1593 | 91.9 | 1.55 | 1203/131.33 | 1196/153.39 |
| 6[e] | 3 | 940 | 871 | 92.8 | 1.71 | 698/34.91 | 649/38.21 |
| 7 | 4 | 1414 | 1344 | 95.2 | 1.68 | 1032/159.16 | 1031/184.82 |
| 8[e] | 4 | 629 | 530 | 84.4 | 1.94 | 490/51.31 | 475/56.22 |
| 9 | 5 | 990 | 728 | 73.6 | 1.44 | 413/141.82 | 406/167.83 |
| 10[e] | 5 | 356 | 332 | 93.5 | 1.80 | 274/39.91 | 259/45.05 |
| 11 | 6 | 1035 | 736 | 71.2 | 1.56 | 421/147.87 | 414/175.86 |
| 12[e] | 6 | 487 | 438 | 90.1 | 1.68 | 357/51.57 | 344/56.42 |
| 13 | 7 | 392 | 149 | 38.0 | 1.56 | 85/204.96 | 85/240.8 |
| 14[e] | 7 | 102 | 69 | 68.3 | 2.35 | 49/95.3 | 48/110.96 |
| 15 | 8 | 468 | 415 | 89.1 | 1.59 | 249/238.03 | 247/276.62 |
| 16[e] | 8 | 195 | 158 | 81.4 | 1.72 | 114/104.2 | 114/114.19 |
| 17 | 9 | 533 | 429 | 80.5 | 1.51 | 206/184.89 | 205/217.32 |
| 18[e] | 9 | 144 | 90 | 62.5 | 1.96 | 64/65.71 | 62/75.13 |
| 19 | 10 | 323 | 305 | 94.7 | 1.58 | 152/23.85 | 65/50.65 |
| 20[e] | 10 | 185 | 160 | 86.5 | 2.00 | 130/87.63 | 127/98.57 |
| 21 | 11 | 758 | 640 | 84.7 | 1.40 | 402/147.81 | 402/174.51 |
| 22[e] | 11 | 148 | 145 | 98.6 | 1.68 | 106/60.44 | 103/70.54 |

[a] $N_{simulations} = 1e11$, SPLIT=11000, $N_{sampled} = 1e8$.

[b] Number of spots found from `dials.find_spots` using spotfinding defaults.

[c] From `dials.index`.

[d] From `dials.tof_integrate`.

[e] With modeled Union beamstop.

Figure S14: Histograms of probabilities for rubredoxin diffraction by pixel for the three detector panels. Note that the color scale is logarithmic. ”box” detector geometry. $V_{crystal} = 1mm^3$. $N_{samples} = 1e11$. $SPLIT =$(a) 1, (b) 10, (c) 100, (d) 1000, (e) 10000.

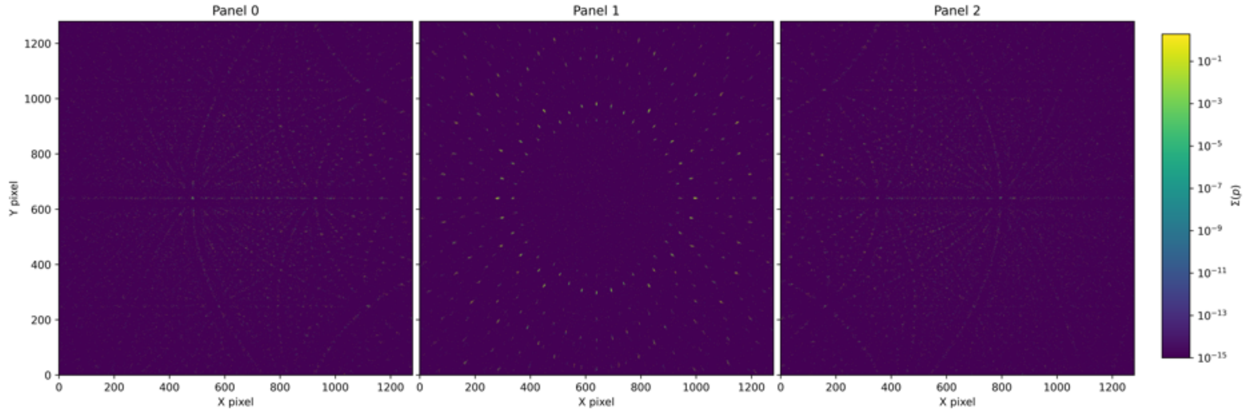


(a)

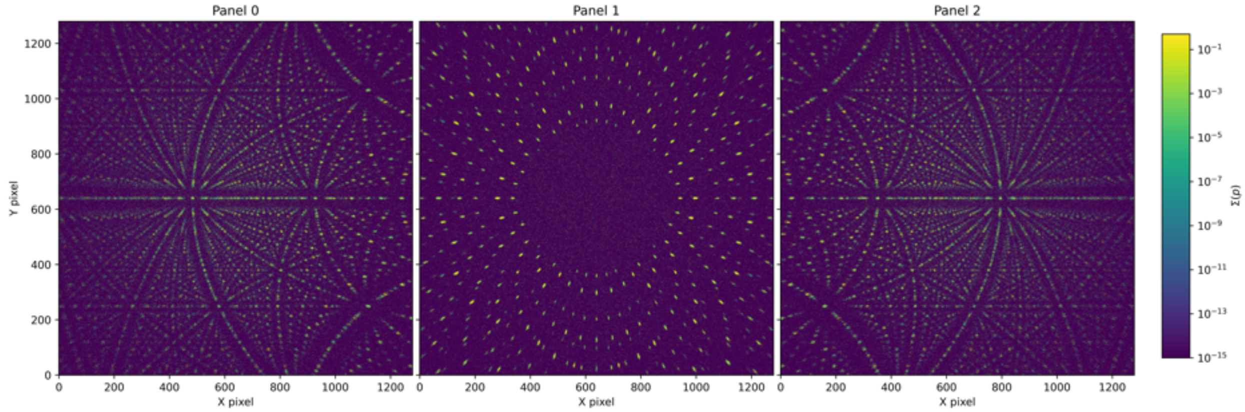


(b)

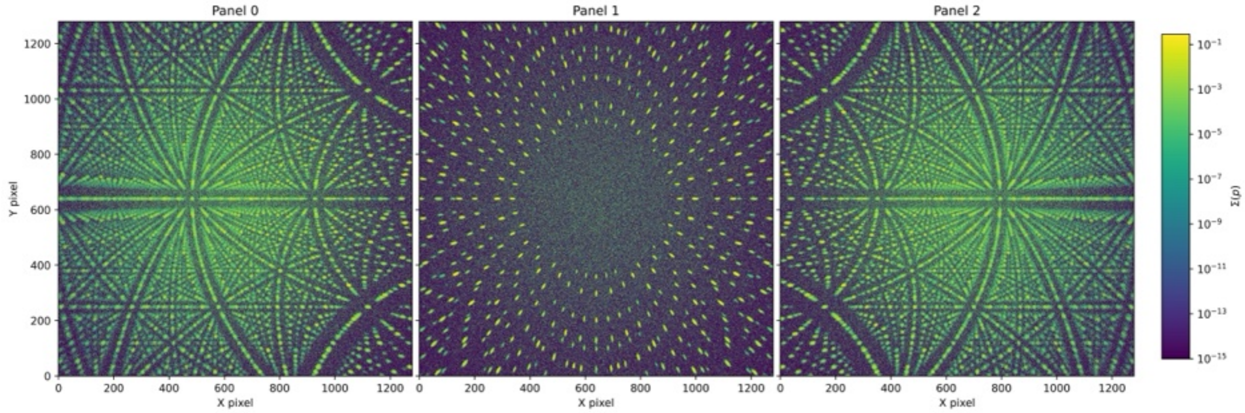


(c)

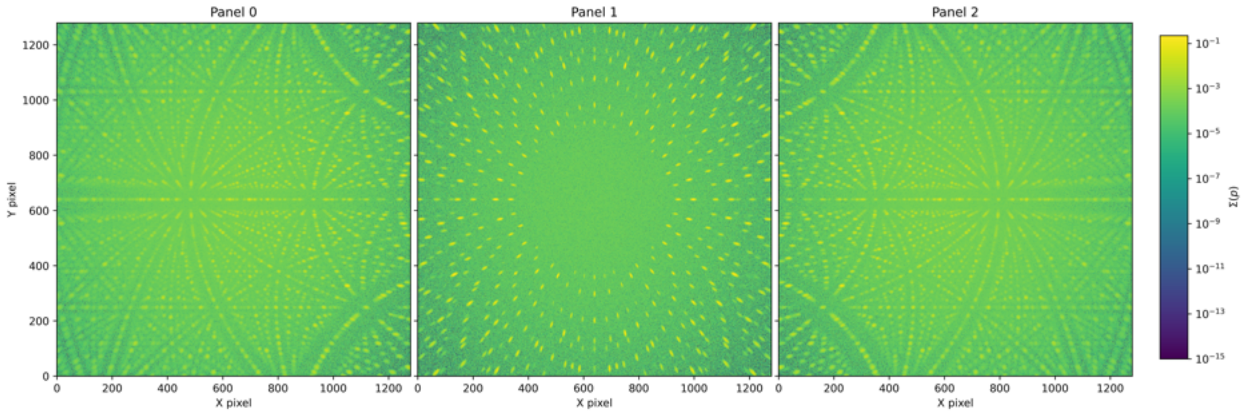
Panel 0
Panel 1
Panel 2
Y pixel
X pixel
Σ(ρ)

(d)

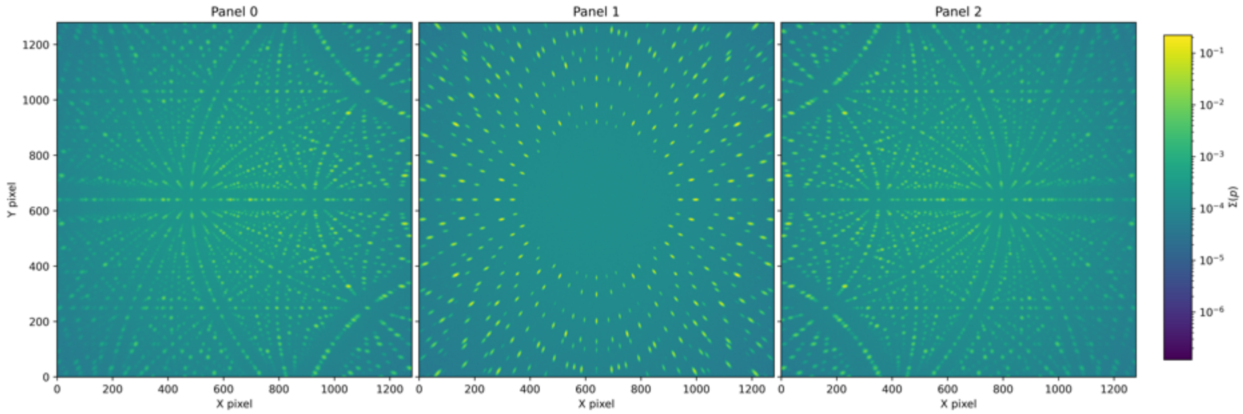
Panel 0
Panel 1
Panel 2
Y pixel
X pixel
Σ(ρ)

(e)
