## Supplementary material for "Simulating Neutron Protein Crystallography Experiments: Applications to the Development of the NMX Instrument at ESS": Calculating Structure factors in Jupyter

calculate\_structure\_factors


### Calculate Structure Factors from PDB Entries for McStas¶

This notebook will take entries from the PDB and calculate the structure factors for use in McStas simulations.

This notebook assumes the pdb data is from neutrons and has structure factor data. If using X-ray data, there are alternative instructions.

#### Why is this necessary?¶

McStas requires a structure factor file in .lau format with a particular formatting. Utilities in McStas for generating .lau files, such as `cif2hkl`, cannot handle the pdb, mtz, and mmcif files from the PDB. Parameters like coherent and incoherent neutron scattering per unit cell volume, as well as neutron absorption, need to be calculated based on the molecular formula and solvent volume. Incoherent scattering in particular is affected by the presence of protons over deuterium. Estimating the effects of H/D substitution is a key target for neutron MX simulations.

The `gemmi` package can calculate the molecular formula based on the atomic coordinate file from the PDB, as well as the solvent volume percentage. The `periodictable` package can calculate the neutron scattering/absorption coefficients.

Even if there exist neutron structure factor data, the reality is that such datasets are rarely fully complete, especially in the highest resolution shells. This is a problem for McStas– if a reflection is not in the .lau file, *it will not be modeled.* (Maybe this is fine! But usually you want all the reflections to be in reciprocal space, at least in theory.) Therefore, calculating all reflections from the atomic coordinates to a particular resolution is required. However, we can use the observed data from the PDB as a starting point for calculating the structure factors' solvent contribution. Since solvent affects low-resolution reflections much more than high-resolution, this is a reasonable assumption to make.

Anomalous contributions are not calculated.

#### What Do I Need?¶

1. The latest Python with Jupyter installed
2. The usual data analysis packages: `numpy`, `scipp`, `pandas`
3. `gemmi` and `periodictable`
4. the CCP4 monomer library (required for gemmi)

#### What does it do?¶

1. Downloads the relevant atomic coordinate and structure factor files from the PDB from an input PDB accession code.
2. Calculates the molecular formula and solvent volume.
3. Calculates the neutron scattering and absorption coefficients.
4. Loads the observed structure factors Fobs from the PDB. Calculates structure factors Fcalc from atomic coordinates with `gemmi`.
5. Generates a solvent mask from the atomic coordinates, and calculates the solvent contribution to structure factors Fsolv.
6. Scales Fcalc and Fsolv with the Fobs data to generate the combined structure factor Ftotal.
7. Expands Ftotal to P1 and calculates all Friedel pairs.
8. Exports all relevant data to .lau format for use in McStas.

In [1]:

```
import h5py
import numpy as np
import scipp as sc
import pandas as pd
import requests
from periodictable import *
import periodictable.fasta
import gemmi
import cmath
from pathlib import Path
from typing import Dict, Mapping
from collections import Counter

import os
os.environ["CLIBD_MON"] = "/Applications/ccp4-9/lib/data/monomers/"
```

### Get Files from the PDB¶

In [2]:

```
"""
Downloads the coordinate, structure factor, and sequence files for
a given PDB accession code.
"""
def download_pdb_files(pdb_id, out_dir="pdb_files") -> Mapping[str, Path | None]:
    pdb_id = pdb_id.lower()
    out_dir = Path(out_dir)
    out_dir.mkdir(exist_ok=True)

    urls = {
        "structure": f"https://files.rcsb.org/download/{pdb_id}.pdb",
        "structure_cif": f"https://files.rcsb.org/download/{pdb_id}.cif",
        "structure_factors": f"https://files.rcsb.org/download/{pdb_id}-sf.cif",  # structure factors
        "fasta": f"https://www.rcsb.org/fasta/entry/{pdb_id}",  # structure factors
    }
    outs = {
        "structure": out_dir / f"{pdb_id}.pdb",
        "structure_cif": out_dir / f"{pdb_id}.cif",
        "structure_factors": out_dir / f"{pdb_id}-sf.cif",  # structure factors
        "fasta": out_dir / f"{pdb_id}.fasta",  # structure factors
    }

    for name, url in urls.items():
        response = requests.get(url)
        if response.status_code == 200:
            filename = outs[name]
            if filename.exists():
                print("Warning: overwriting... ",end="")
            with open(filename, "wb") as f:
                f.write(response.content)
            print(f"Downloaded {name} to {filename}")
        else:
            outs[name] = None
            print(f"Failed to download {name} from {url}")
    return outs

"""
If the files from the PDB already exist, find them
"""
def get_pdb_files_local(pdb_id:str, indir:Path|str) -> Dict[str,Path|None] | None:
    if isinstance(indir, str):
        indir = Path(indir)
    if not indir.exists():
        print(f"{str(indir)} not found")
        return None
    outs = {}
    structure = indir  / f"{pdb_id}.pdb"
    if structure.exists():
        outs["structure"] = structure
    else:
        print(f"Structure file {str(structure)} not found.")
        outs["structure"] = None

    structure_cif = indir / f"{pdb_id}.cif"
    if structure_cif.exists():
        outs["structure_cif"] = structure_cif
    else:
        print(f"structure_cif file {str(structure_cif)} not found.")
        outs["structure_cif"] = None

    structure_factors = indir / f"{pdb_id}-sf.cif"
    if structure_factors.exists():
        outs["structure_factors"] = structure_factors
    else:
        print(f"structure_factors file {str(structure_factors)} not found.")
        outs["structure_factors"] = None

    fasta = indir / f"{pdb_id}.fasta"
    if fasta.exists():
        outs["fasta"] = fasta
    else:
        print(f"fasta file {str(fasta)} not found.")
        outs["fasta"] = None

    return outs
```

In [ ]:

```
# Download the PDB files
pdb_files = download_pdb_files("4AR4", "4AR4")

# If you already have the PDB files
# pdb_files = get_pdb_files_local(
#     "4ar4", Path("/Users/aaronfinke/calculated_structure_factors/4ar4")
# )
```

In [ ]:

```
structure_name = "rubredoxin"
parent_dir = pdb_files["structure"].parent
```

In [7]:

```
st = gemmi.read_structure(str(pdb_files['structure']))
st.setup_entities()
for ent in st.entities:
    print(ent.name)
    print(ent.subchains)
    print(ent.entity_type)
    print(ent.polymer_type)
    print(type(gemmi.PolymerType.PeptideL))
    if ent.polymer_type == gemmi.PolymerType.PeptideL:
        print(gemmi.one_letter_code(ent.full_sequence))
```

```
A
['Axp']
EntityType.Polymer
PolymerType.PeptideL
<enum 'PolymerType'>
MAKWVCKICGYIYDEDAGDPDNGISPGTKFEELPDDWVCPICGAPKSEFEKLED
FE!
['Ax1']
EntityType.NonPolymer
PolymerType.Unknown
<enum 'PolymerType'>
D8U!
['Ax2', 'Ax3']
EntityType.NonPolymer
PolymerType.Unknown
<enum 'PolymerType'>
D3O!
['Ax4', 'Ax5', 'Ax6', 'Ax7']
EntityType.NonPolymer
PolymerType.Unknown
<enum 'PolymerType'>
water
['Axw']
EntityType.Water
PolymerType.Unknown
<enum 'PolymerType'>
```

In [73]:

```
output_dir = str(pdb_files["structure_factors"].parent)
ciffile = str(pdb_files["structure_factors"])
mtzfile = str(pdb_files["structure_factors"].with_suffix('.mtz'))

# generate the mtz file from the structure factor cif file
# uses the command line program as there is no pythonic alternative to do this
! gemmi cif2mtz {ciffile} {mtzfile}
pdb_files["structure_factors_mtz"] = Path(mtzfile)
```

In [5]:

```
# get fasta sequence
with open(pdb_files["fasta"]) as fp:
# with open("/Users/aaronfinke/calculated_structure_factors/3ryg.xyz") as fp:
    fasta_str = fp.read()
fasta_seq = gemmi.read_pir_or_fasta(fasta_str)
fasta_seq[0].seq
```

Out[5]:

```
'MAKWVCKICGYIYDEDAGDPDNGISPGTKFEELPDDWVCPICGAPKSEFEKLED'
```

In [6]:

```
# get molecular formula from fasta
seq = periodictable.fasta.Sequence("seq", fasta_seq[0].seq, type="aa")

print(seq.formula)
```

```
C267H309H[1]75N62O87S5
```

In [7]:

```
#get molecular formula from pdb file
#note this includes all defined waters and ligands, but that's fine
import gemmi 
st = gemmi.read_structure(str(pdb_files["structure"]))
# st.remove_alternative_conformations()
element_counts = Counter()

# count elements
for model in st:
    for chain in model:
        for residue in chain:
                for atom in residue:
                    if atom.element != gemmi.Element('X'):
                        element_counts[str(atom.element.name)] += atom.occ
print(f"{element_counts=}")
element_counts
```

```
element_counts=Counter({'D': 496.8300006836653, 'C': 267.0000000298023, 'O': 175.36999996751547, 'N': 62.00000002980232, 'H': 6.480000011622906, 'S': 5.0, 'Fe': 1.0})
```

Out[7]:

```
Counter({'D': 496.8300006836653,
         'C': 267.0000000298023,
         'O': 175.36999996751547,
         'N': 62.00000002980232,
         'H': 6.480000011622906,
         'S': 5.0,
         'Fe': 1.0})
```

In [8]:

```
# turn molecular formula from pdb file into string for periodictable
for k,v in element_counts.items():
    if round(v,3).is_integer():
        element_counts[k] = int(v)

element_string = {key:str(round(value,2)) for key, value in element_counts.items() }
for k,v in element_string.items():
    if v == '1':
        element_string[k] = ''


formula_string = " ".join(f"{k}{round(v,2)}" for k,v in element_counts.items())
formatted_formula_string = " ".join(
    f"{k}{v}" for k, v in element_string.items()
)
formula_string, formatted_formula_string
```

Out[8]:

```
('N62 C267 O175.37 S5 D496.83 H6.48 Fe1',
 'N62 C267 O175.37 S5 D496.83 H6.48 Fe')
```

In [9]:

```
# Calculate solvent content
protein_density = 1.35      # density of protein in g/mL; usually around 1.34-1.35

# if there is only one AA chain
st = gemmi.read_structure(str(pdb_files["structure"]))
st.setup_entities()
chain = st[0]["A"]
poly = chain.get_polymer()
ent = st.get_entity_of(poly)
sequence = ent.full_sequence
weight = gemmi.calculate_sequence_weight(ent.full_sequence)
print(f"Molecular weight of protein: {weight:.2f}")
matthews_coeff = st.cell.volume_per_image() / weight
print(f"Matthews coefficient: {matthews_coeff:.2f}")
protein_fraction = 1.0 / (6.02214e23 * 1e-24 * protein_density * matthews_coeff)
print("Solvent content: {:.1f}%".format(100 * (1 - protein_fraction)))
solv_cont = f"{100 * (1 - protein_fraction):.1f}"
protein_cont = round(100 * protein_fraction,1)
```

```
Molecular weight of protein: 6031.73
Matthews coefficient: 2.21
Solvent content: 44.2%
```

In [10]:

```
# Calculate solvent content

# multiple AA chains
weight=0
st = gemmi.read_structure(str(pdb_files["structure"]))
st.setup_entities()
for model in st:
    for chain in model:
        poly = chain.get_polymer()
        ent = st.get_entity_of(poly)
        sequence = ent.full_sequence
        subweight = gemmi.calculate_sequence_weight(ent.full_sequence)
        weight += subweight

matthews_coeff = st.cell.volume_per_image() / weight
protein_fraction = 1.0 / (6.02214e23 * 1e-24 * protein_density * matthews_coeff)
solv_cont = f"{100 * (1 - protein_fraction):.1f}"
protein_cont = round(100 * protein_fraction, 1)

print(f"Matthews coeff: {matthews_coeff:0.2f}")
print("Solvent content: {:.1f}%".format(100 * (1 - protein_fraction)))
```

```
Matthews coeff: 2.21
Solvent content: 44.2%
```

In [13]:

```
# replace all H with D

# uncomment if you want to do this

# mon_lib = gemmi.read_monomer_lib(os.environ["CLIBD_MON"], st[0].get_all_residue_names())

# topo = gemmi.prepare_topology(st=st, monlib=mon_lib, model_index=0, h_change=gemmi.HydrogenChange.ReAdd)

# for chain_info in topo.chain_infos:
#     for res_info in chain_info.res_infos:
#         res = res_info.res
#         for atom in res:
#             if atom.element == gemmi.Element("H"):
#                 atom.name = atom.name.replace("H", "D")
#                 atom.element = gemmi.Element("D")
```

#### Calculate neutron scattering and absorption cross-sections¶

In [ ]:

```
sigma_coh = 0
sigma_inc = 0
sigma_abs = 0

seq_cryst = periodictable.formula(
    f"{protein_cont}%vol (100%wt {formula_string}@{protein_density}) // D2O@1n"
)
for el, amt in seq_cryst.atoms.items():
    sigma_coh += el.neutron.coherent * amt
    incoh = el.neutron.incoherent
    sigma_inc += incoh * amt
    sigma_abs += el.neutron.absorption

print(f"Coherent scattering cross-section: {sigma_coh}") 
print(f"Incoherent scattering cross-section: {sigma_inc}") 
print(f"Absorption cross-section: {sigma_abs}")
```

```
Coherent scattering cross-section: 9782.136157428069
Incoherent scattering cross-section: 2652.5390695287424
Absorption cross-section: 188.16976523785098
```

#### Generate the calculated structure factors¶

In [81]:

```
# read observed structure factors from the PDB

mtz = gemmi.read_mtz_file(str(pdb_files["structure_factors_mtz"]))
mtz.update_reso()
f_obs = mtz.get_value_sigma("FP", "SIGFP")
```

In [76]:

```
# summation SF calculation
d_min = 1.0

st = gemmi.read_structure(str(pdb_files["structure"]))
cell = st.cell
spacegroup = gemmi.SpaceGroup(st.spacegroup_hm)
dc = gemmi.StructureFactorCalculatorN(cell)
gops = spacegroup.operations()
hkl_f = []
refl_array = gemmi.make_miller_array(st.cell, spacegroup, d_min)
for refl in refl_array:
    if not gops.is_systematically_absent(refl):
        x = dc.calculate_sf_from_model(st[0], refl)
        f = abs(x)
        f2 = f ** 2
        hkl_f.append((refl,x,round(f,2),round(f2,2)))
```

In [ ]:

```
mtz = gemmi.Mtz(with_base=True)
mtz.set_cell_for_all(st.cell)
mtz.spacegroup = st.find_spacegroup()
mtz.add_dataset("calculated")
mtz.add_column("FC", "F", -1, -1)
mtz.add_column("IC", "I", -1, -1)
mtz.add_column("PHIC", "P", -1, -1)
print(mtz.column_labels())
mtz.set_data(
    np.array(
        [
            [
                hkl[0],
                hkl[1],
                hkl[2],
                f,
                f2,
                cmath.phase(x),
            ]
            for hkl, x, f, f2 in hkl_f
        ]
    )
)

mtz.expand_to_p1()
```

Out[ ]:

```
<gemmi.Mtz with 4 columns, 515711 reflections>
```

In [ ]:

```
# Now add Friedel pairs...
mtz_df = pd.DataFrame(data=mtz.array, columns=mtz.column_labels())
mtz_df = mtz_df.astype({label: "int32" for label in "HKL"})


neg_df = mtz_df.copy()
neg_df[["H", "K", "L"]] = -neg_df[["H", "K", "L"]]
df_extended = pd.concat([mtz_df, neg_df], ignore_index=True)
```

Out[ ]:

|  | H | K | L | IMODEL |
| --- | --- | --- | --- | --- |
| 0 | 0 | 0 | 2 | 1.893166e+06 |
| 1 | 0 | 0 | 4 | 2.032938e+06 |
| 2 | 0 | 0 | 6 | 1.047465e+05 |
| 3 | 0 | 0 | 8 | 2.036476e+05 |
| 4 | 0 | 0 | 10 | 4.412282e+05 |
| ... | ... | ... | ... | ... |
| 1031417 | -57 | 5 | 2 | 2.111637e-02 |
| 1031418 | 57 | -5 | 2 | 2.111637e-02 |
| 1031419 | 57 | 5 | -3 | 1.261715e-01 |
| 1031420 | -57 | 5 | 3 | 1.261715e-01 |
| 1031421 | 57 | -5 | 3 | 1.261715e-01 |

1031422 rows × 4 columns

In [ ]:

```
# write the McStas .lau file

sg = st.find_spacegroup()
uc = st.cell
with open(
    parent_dir
    / f"{''.join([x if x.isalnum() else '_' for x in structure_name])}.lau",
    "w",
) as fp:
    fp.write(
        f"# {structure_name.capitalize()} (F^2calc), {formatted_formula_string}'\n"
    )
    fp.write(f"# SPCGRP {sg.hm} ({sg.ccp4}) {sg.crystal_system().name}\n")
    fp.write(
        f"# a={uc.a:.2f}, b={uc.b:.2f}, c={uc.c:.2f}; alpha={uc.alpha:.1f}, beta={uc.beta:.1f}, gamma={uc.gamma:.1f};\n"
    )
    fp.write("# Reference: (TO BE ADDED)\n")
    fp.write("\n")
    fp.write("# Physical parameters:\n")
    fp.write(f"# lattice_a {uc.a}   lattice parameter a in [Angs]\n")
    fp.write(f"# lattice_b {uc.b}   lattice parameter b in [Angs]\n")
    fp.write(f"# lattice_c {uc.c}  lattice parameter c in [Angs]\n")
    fp.write(f"# lattice_aa {uc.alpha}     lattice angle alpha in [deg]\n")
    fp.write(f"# lattice_bb {uc.beta}     lattice angle alpha in [deg]\n")
    fp.write(f"# lattice_cc {uc.gamma}     lattice angle alpha in [deg]\n")
    fp.write(f"# weight  {weight:.3f}   in [g/mol] (single protein)\n")
    fp.write(
        f"# sigma_coh {sigma_coh:.1f}  coherent scattering cross section (protein+solvent) in [barn]\n"
    )
    fp.write(
        f"# sigma_abs {sigma_abs:.1f} absorption scattering cross section (protein+solvent) in [barn]\n"
    )
    fp.write(
        f"# sigma_inc {sigma_inc:.1f}  incoherent scattering cross section (protein+solvent) in [barn]\n"
    )
    fp.write("#\n")
    fp.write("# Format parameters: Crystallographica format\n")
    fp.write("# column_F2 4 norm of scattering factor |F|^2 in [fm^2]\n")
    fp.write("# column_h 1\n")
    fp.write("# column_k 2\n")
    fp.write("# column_l 3\n")
    fp.write("\n")

    for index, row in df_extended.iterrows():
        
        print(
            f"{int(row['H'])} {int(row['K'])} {int(row['L'])} {row['IMODEL']:.2f}",
            file=fp,
        )
```

#### Generate Structure Factors with Solvent Density¶

It is recommended to do this with `phenix.fmodel` but you can also use Gemmi, as shown here.

In [105]:

```
# calculate ideal structure factors from the atomic coordinates
d_min = 1.0

st = gemmi.read_structure(
    str(pdb_files['structure'])
)
d_min =1.0 - 1e-9
dc = gemmi.DensityCalculatorN()
dc.d_min = d_min
dc.set_refmac_compatible_blur(st[0])
dc.grid.setup_from(st)
dc.put_model_density_on_grid(st[0])
grid = gemmi.transform_map_to_f_phi(dc.grid, half_l=True)
f_cryst = grid.prepare_asu_data(dmin=d_min, unblur=dc.blur)
```

In [17]:

```
#prepare solvent mask

mask_grid = gemmi.FloatGrid()
mask_grid.setup_from(st, spacing=min(0.6, d_min / 2 - 1e-9))
masker = gemmi.SolventMasker(gemmi.AtomicRadiiSet.Refmac)
masker.put_mask_on_float_grid(mask_grid, st[0])
fmask_gr = gemmi.transform_map_to_f_phi(mask_grid, half_l=True)
f_mask = fmask_gr.prepare_asu_data(dmin=d_min)
len(f_mask), len(f_cryst), len(f_obs)
```

Out[17]:

```
(14520, 14520, 11122)
```

In [18]:

```
#scale solvent mask with calc'd / observed F
scaling = gemmi.Scaling(st.cell, st.find_spacegroup())
scaling.use_solvent = True
scaling.prepare_points(f_cryst, f_obs, f_mask)
scaling.fit_isotropic_b_approximately()
wssr = scaling.fit_parameters()
# print(f"RMSE: {(wssr / len(f_obs)) ** 0.5:.4f}")
print(f"R-factor: {scaling.calculate_r_factor():.2%}")
```

```
R-factor: 21.41%
```

In [19]:

```
f_tot = f_cryst.copy()
scaling.scale_data(f_tot, f_mask)
```

In [83]:

```
"""Get phase of complex HKL value and convert to degrees"""
def phase_in_angles(v: gemmi.ComplexHklValue, eps: float = 2e-5) -> float:
    angle = np.rad2deg(cmath.phase(v.value))
    if angle < -eps:
        angle += 360.0
    return max(0.0, angle)
```

In [21]:

```
# setup gemmi mtz with symmetry

mtz = gemmi.Mtz(with_base=True)
mtz.set_cell_for_all(st.cell)
mtz.spacegroup = st.find_spacegroup()
mtz.add_dataset("calculated")
mtz.add_column("FC", "F", -1, -1)
mtz.add_column("IC", "I", -1, -1)
mtz.add_column("PHIC", "P", -1, -1)
print(mtz.column_labels())
mtz.set_data(
    np.array(
        [
            [
                f_cryst[i].hkl[0],
                f_cryst[i].hkl[1],
                f_cryst[i].hkl[2],
                abs(f_cryst[i].value),
                abs(f_cryst[i].value) ** 2,
                phase_in_angles(f_cryst[i]),
            ]
            for i in range(len(f_cryst))
        ]
    )
)
mtz.expand_to_p1()
```

```
['H', 'K', 'L', 'FC', 'IC', 'PHIC']
```

In [22]:

```
# Now add Friedel pairs...
mtz_df = pd.DataFrame(data=mtz.array, columns=mtz.column_labels())
mtz_df = mtz_df.astype({label: "int32" for label in "HKL"})


neg_df = mtz_df.copy()
neg_df[["H", "K", "L"]] = -neg_df[["H", "K", "L"]]
df_extended = pd.concat([mtz_df, neg_df], ignore_index=True)
df_extended
```

Out[22]:

|  | H | K | L | FC | IC | PHIC |
| --- | --- | --- | --- | --- | --- | --- |
| 0 | -20 | 0 | 1 | 92.203529 | 8501.491211 | 180.000015 |
| 1 | -20 | 0 | 2 | 42.842777 | 1835.503540 | 0.000000 |
| 2 | -20 | 0 | 3 | 43.216034 | 1867.625610 | 0.000000 |
| 3 | -20 | 0 | 4 | 36.677803 | 1345.261230 | 180.000015 |
| 4 | -20 | 0 | 5 | 123.369759 | 15220.097656 | 0.000019 |
| ... | ... | ... | ... | ... | ... | ... |
| 55625 | 20 | -4 | 0 | 26.897518 | 723.476440 | 145.806564 |
| 55626 | 20 | -4 | 1 | 62.623798 | 3921.740234 | 77.386505 |
| 55627 | 20 | -4 | 2 | 40.156387 | 1612.535522 | 319.627716 |
| 55628 | 20 | -4 | 3 | 44.795189 | 2006.608887 | 207.249771 |
| 55629 | 20 | -4 | 4 | 12.119370 | 146.879150 | 225.849670 |

55630 rows × 6 columns

#### Save the reflections as a .lau file for McStas¶

In [92]:

```
sg = st.find_spacegroup()
uc = st.cell
with open(
    parent_dir / f"{''.join([x if x.isalnum() else '_' for x in structure_name])}.lau", "w"
) as fp:
    fp.write(
        f"# {structure_name.capitalize()} (F^2calc), {formatted_formula_string}'\n"
    )
    fp.write(f"# SPCGRP {sg.hm} ({sg.ccp4}) {sg.crystal_system().name}\n")
    fp.write(
        f"# a={uc.a:.2f}, b={uc.b:.2f}, c={uc.c:.2f}; alpha={uc.alpha:.1f}, beta={uc.beta:.1f}, gamma={uc.gamma:.1f};\n"
    )
    fp.write("# Reference: (TO BE ADDED)\n")
    fp.write("\n")
    fp.write("# Physical parameters:\n")
    fp.write(f"# lattice_a {uc.a}   lattice parameter a in [Angs]\n")
    fp.write(f"# lattice_b {uc.b}   lattice parameter b in [Angs]\n")
    fp.write(f"# lattice_c {uc.c}  lattice parameter c in [Angs]\n")
    fp.write(f"# lattice_aa {uc.alpha}     lattice angle alpha in [deg]\n")
    fp.write(f"# lattice_bb {uc.beta}     lattice angle alpha in [deg]\n")
    fp.write(f"# lattice_cc {uc.gamma}     lattice angle alpha in [deg]\n")
    fp.write(f"# weight  {weight:.3f}   in [g/mol] (single protein)\n")
    fp.write(
        f"# sigma_coh {sigma_coh:.1f}  coherent scattering cross section (protein+solvent) in [barn]\n"
    )
    fp.write(
        f"# sigma_abs {sigma_abs:.1f} absorption scattering cross section (protein+solvent) in [barn]\n"
    )
    fp.write(
        f"# sigma_inc {sigma_inc:.1f}  incoherent scattering cross section (protein+solvent) in [barn]\n"
    )
    fp.write("#\n")
    fp.write("# Format parameters: Crystallographica format\n")
    fp.write("# column_F2 4 norm of scattering factor |F|^2 in [fm^2]\n")
    fp.write("# column_h 1\n")
    fp.write("# column_k 2\n")
    fp.write("# column_l 3\n")
    fp.write("\n")

    for index, row in df_extended.iterrows():
        print(
            f"{int(row['H'])} {int(row['K'])} {int(row['L'])} {row['IC']:.6f}",
            file=fp,
        )
```
